# Characterizing organisms from three domains of life with universal primers from throughout the global ocean

**DOI:** 10.1101/2025.02.19.638942

**Authors:** Jesse McNichol, Nathan L R Williams, Yubin Raut, Craig Carlson, Elisa R Halewood, Kendra Turk-Kubo, Jonathan P Zehr, Andrew P Rees, Glen Tarran, Mary R. Gradoville, Matthias Wietz, Christina Bienhold, Katja Metfies, Sinhué Torres-Valdés, Thomas Mock, Sarah Lena Eggers, Wade Jeffrey, Joseph Moss, Paul Berube, Steven Biller, Levente Bodrossy, Jodie Van De Kamp, Mark Brown, Swan L. S. Sow, E. Virginia Armbrust, Jed Fuhrman

## Abstract

We introduce the Global rRNA Universal Metabarcoding Plankton database (GRUMP), which consists of 1194 samples that were collected from 2003-2020 and cover extensive latitudinal and longitudinal transects, as well as depth profiles in all major ocean basins. DNA from unfractionated (> 0.2µm) seawater samples was amplified using the 515Y/926R universal three- domain rRNA gene primers, simultaneously quantifying the relative abundance of amplicon sequencing variants (ASVs) from bacteria, archaea, eukaryotic nuclear 18S, and eukaryotic plastid 16S. Thus, the ratio between taxa in one sample is directly comparable to the ratio in any other GRUMP sample, regardless of gene copy number differences. This obviates a problem in prior global studies that used size-fractionation and different rRNA gene primers for bacteria, archaea, and eukaryotes, precluding comparisons across size fractions or domains. On average, bacteria contributed 71%, eukaryotes 19%, and archaea 8% to rRNA gene abundance, though eukaryotes contributed 32% at latitudes > 40°. GRUMP is publicly available on the Simons Collaborative Marine Atlas Project (CMAP), promoting the global comparison of marine microbial dynamics.

## 1.0 Background & Summary / Introduction

Microorganisms span all three domains of life: Archaea, Bacteria, and Eukarya^1^. Together, these microbes are critical to life on Earth. Photosynthetic marine cyanobacteria and their eukaryotic counterparts, collectively called phytoplankton, form the base of marine food webs, thereby sustaining productivity and biodiversity^2–5^. The bacterioplankton, which consist of bacteria and archaea, play critical roles in biogeochemical cycles: they consume organic matter and biogeochemically transform and remineralize inorganic and organic compounds, mediating the magnitude of carbon sequestered in the deep sea^5^. Additionally, heterotrophic protists and larger zooplankton impose a top-down control on the plankton community via grazing, with concomitant respiration. To understand the fundamental ecological question of where these microbes are and why, it is vital to study how biological interactions structure plankton communities^6,7^, which calls for direct quantitative comparisons. So far, only a handful of studies in oceanography have used metabarcoding techniques that allow the direct and quantitative comparison of organisms from all three domains of life^8–10^, limiting the field of marine microbial ecology and biological oceanography to a fragmented evaluation of the ocean’s microbial and biogeochemical processes.

The emergence of high-throughput sequencing has significantly enhanced our ability to robustly identify microbial taxa. However, many current metagenomic or metabarcoding datasets that focus on the global distribution of oceanic microbes do not allow direct quantitative comparisons across all three domains of life. Studies such as TARA Oceans^13^, Malaspina^14^, and KAUST Metagenomic Analysis Platform (KMAP)^15^ have provided key insights into microbial distributions and drivers, yet they generally use different operationally defined size fractions to separate smaller from larger organisms. This precludes comparisons between samples that have been quantified compositionally rather than in absolute units. Other marine metagenomic studies such as bioGEOTRACES^16^ and Bio-GO-SHIP^17^ have used unfractionated seawater samples, but these metagenome-based studies remain limited by our inability to fully classify metagenomic reads^15^, meaning that a complete community profile is difficult to obtain. Furthermore, many studies also use separate primer pairs to amplify eukaryotic 18S rRNA and prokaryotic 16S rRNA for community analysis from whole seawater samples, yielding separate compositional data on each^18,19^. Whilst these methods have been effective at characterizing the microbial communities separately, they prevent the direct, holistic comparison between eukaryotes, bacteria, and archaea.

To address this shortcoming, we utilized unfractionated samples and applied a universal primer pair (515Y/926R) that amplifies bacteria, archaea, and eukaryotes; thus allowing for the quantitative characterization of the entire microbial community in a single PCR reaction^20^. The resulting 16S and 18S rRNA sequences are denoised into amplicon sequence variants (ASVs) using DADA2 in QIIME2^21,22^, providing resolution down to single-nucleotide differences. This primer pair has been validated as robust and highly accurate via both mock communities of marine bacteria, archaea^20^, and eukaryotes^23^; such accuracy is not observed with other commonly tested rRNA gene primers^23,24^. Furthermore, comparisons with epipelagic metagenomes from around the world have shown that these universal rRNA gene primers perfectly match 96% of the corresponding genomic primer regions across all three domains^24^. Most importantly, this means that the relative abundance of amplicon sequencing variants (ASVs) from bacteria, archaea, and eukaryotes are able to be quantified together as a total community. We used the 515Y/926R primer pair to sequence the microbial community from 1194 non-fractionated seawater samples from the Arctic, Atlantic, Pacific, Indian, and Southern Oceans. Samples were taken on 19 cruise transects that spanned all four seasons, including cruises with similar spatial coverage but in different years and seasons. Multiple transects also have high depth coverage, ranging from the surface to 2000 m, and in some cases to just above the ocean floor (> 6000 m). Collectively, this database, the **G**lobal **r**RNA **U**niversal **M**etabarcoding of **P**lankton database, or GRUMP, provides coverage of the marine microbial community across diverse ocean biomes and spatiotemporal scales. We have made this dataset publicly available on the Simons Foundation Collaborative Marine Atlas Project CMAP, as well as on the Open Science Framework, OSF representing a powerful tool to compare distributions of marine microbes at a global scale.

## Methods

### Sample Collection and Data Compilation

Samples were collected through multiple collaborations (for exact locations see Figure 1), each of which employed slightly different sampling techniques. For the Atlantic Meridional Transects (AMT 19 and AMT 20), 5 – 10 L of whole seawater were collected from the surface using a Niskin bottle and filtered onto 0.22 µm Sterivex filters. Sterivex filters were capped and stored in RNAlater at -80°C until DNA extraction.

**Figure 1.**
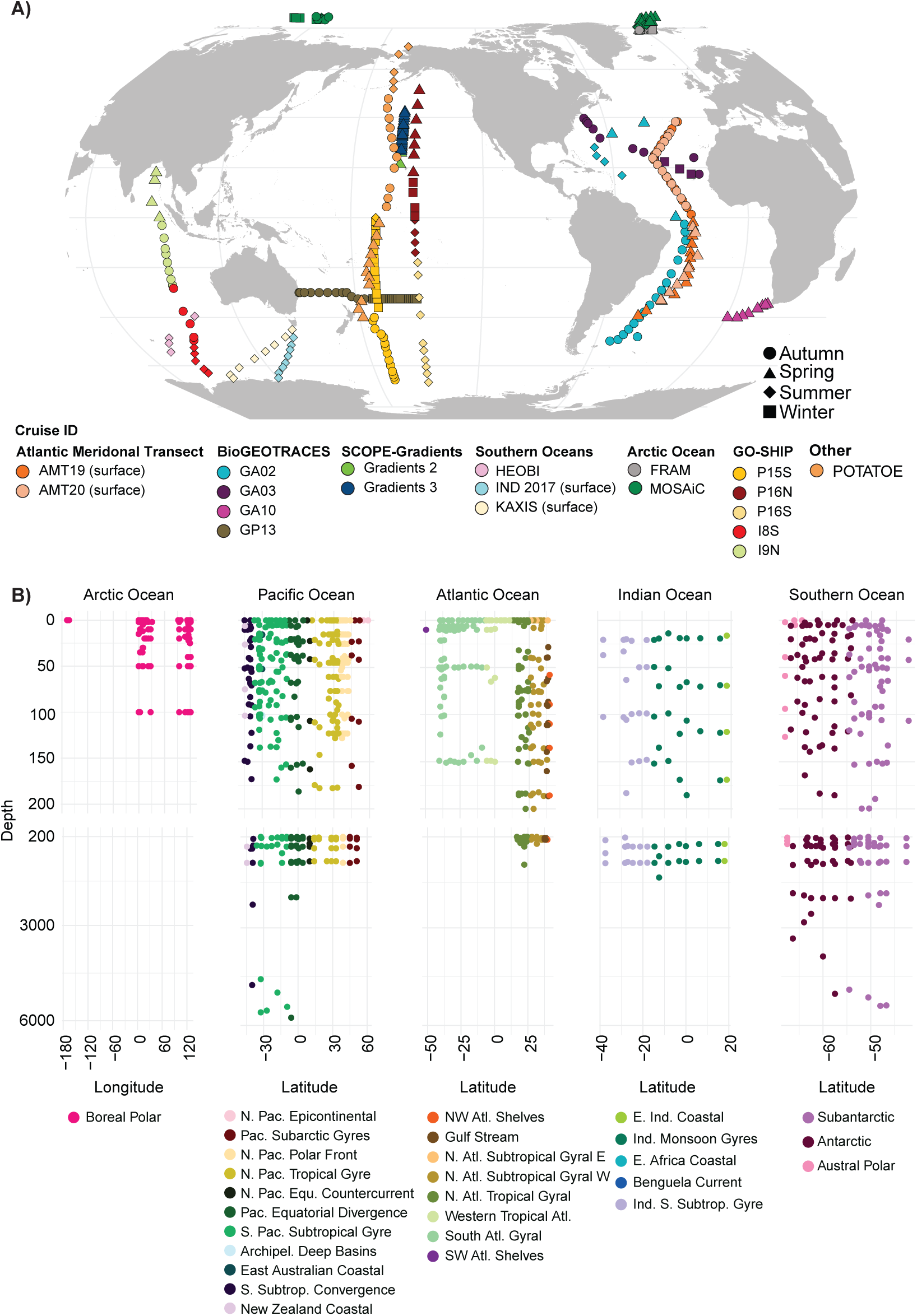
a. Map of the locations of samples in the GRUMP dataset. Colour indicates the cruise names. Shape indicates the season in which the sample was taken. **b.** Depth profile within each ocean basin. Color refers to the Longhurst province of each sample location.

For FRAM, whole seawater was autonomously collected using Remote Access Samplers (RAS; McLane) on four seafloor moorings F4-S-1, HG-IV-S-1, Fevi-34, and EGC-5 (Supplementary Figure 1) from July 2016 – August 2017 in ∼biweekly intervals. Moorings were operated within the FRAM/ Hausgarten Observatory covering the West Spitsbergen Current, central Fram Strait, and East Greenland Current, with nominal deployment depths of 30 m (F4, HG-IV), 67 m (Fevi), and 80 m (EGC). Vertical movements in the water column resulted in variability in the actual sampling depths, ranging from 25 - 150 m. Per sampling event, approximately 700 mL of mercuric-chloride fixed water was filtered onto 0.22 µm Sterivex filters, which were stored at - 20°C until DNA extraction.

For Multidisciplinary drifting Observatory for the Study of Arctic Climate (MOSAiC), whole seawater was collected from the upper water column via a rosette sampler equipped with Niskin bottles deployed through a hole in the sea ice next to the RV *Polarstern*. Where possible, two Niskin bottles per depth were used to collect duplicate samples during the up-casts, nominally at the surface (∼5 m), 10 m, chlorophyll maximum (∼20 – 40 m), 50 m, and 100 m. From these Niskin bottles, 1 - 4 L of whole seawater were filtered onto 0.22 µm Sterivex filters in a temperature-controlled lab at 1°C illuminated with only red light. The number of Sterivex filters per sample varied between two during polar night and 3 - 4 during polar day, depending on the biomass load of the samples. Sterivex filters were stored at -80°C until DNA extraction.

BioGEOTRACES cruises, including transects GA02, GA03, GA10, and GP13, collected whole seawater using a Niskin bottle from between the surface - 5600 m, filtering 100 mL of seawater onto 0.2 µm, 25 mm polycarbonate filters. After filtration, 3 mL of sterile preservation solution (10 mM Tris, pH 8.0; 100 mM EDTA; 0.5 M NaCl) was added, and samples were stored in cryovials at -80°C until DNA extraction.

During the 2017 and 2019 SCOPE (Simons Collaboration on Ocean Processes and Ecology) - Gradients cruises on the R/V Marcus and the R/V Kilo Moana respectively, 0.7 – 4 L of whole seawater was collected using the ship’s underway system, which is approximately 7 m below the surface, as well as the CTD rosette sampler for depths between 15 – 125 m. This water was filtered onto 0.22 µm, 25 mm Supor membrane filters and stored at -80°C until DNA extraction.

The collection of the Southern Ocean transects including the 1) IND-2017 dataset from the Totten Glacier-Sabrina Coast voyage in 2017, 2) the Kerguelen-Axis Marine Science program (K-AXIS) on the Australian Antarctic Division RV *Aurora Australis* Voyage 3 in 2015 - 2016, 3) the Global Ocean Ship-based Hydrographic Investigations Program (GO-SHIP) P15S cruise in 2016, and 4) the Heard Earth-Ocean-Biosphere Interactions (HEOBI) voyage in 2016. For these cruises, 2 L of whole seawater was filtered onto 0.22 µm Sterivex filters. This water was collected from the ships underway during IND-2017, from between 5 - 4625 m during K-AXIS, from between 5 - 6015 m during GO-SHIP P15S, and from between 7 - 3579 m during HEOBI. After filtration, samples were stored at -80°C until DNA extraction.

As part of US GO-SHIP, the additional transects I08S, I09N, P16S, and P16N included some that also traversed into the Southern Ocean (i.e., I08S, P16S) or the Arctic Ocean (P16N). For I08S and I09N, 2 L of whole seawater was collected via Niskin bottle from between 5 – 1450 m depth and was then filtered onto 0.22 µm, 25 mm Supor membrane filters during the cruises in 2007 aboard the R/V *Roger Revelle*. Sucrose lysis buffer was added to the filters, which were then stored at -80°C until DNA extraction. For P16N, whole seawater was collected via niskin bottle from between 20 – 1000 m and for P16S, between 2 – 500 m. Seawater was filtered onto 0.22 µm, 25 mm Supor filters during two latitudinal transects of the Pacific Ocean in 2005 and 2006 as part of the US GO-SHIP repeat hydrography program (previously CLIVAR). Samples were stored as partially extracted lysates in sucrose lysis buffer at -80°C until DNA extraction.

Finally, for samples from the Production Observations Through Another Trans-Latitudinal Oceanic Expedition (POTATOE) cruise, 20 L of whole seawater was collected from the sea surface between 1 – 2 m via bucket casting or Go-Flo bottles and filtered onto 0.22 µm Sterivex filters during a “ship of opportunity” cruise on the RVIB *Nathaniel B Palmer* in 2003^25,26^. Sterivex filters were stored dry at -80°C until DNA extraction.

All datasets have corresponding environmental metadata. We included date, time, latitude, longitude, depth, temperature, salinity, oxygen for all transects, and nutrient data where available. However, some cruises have other environmental data, which can be found at the British Oceanographic Data Centre (https://www.bodc.ac.uk/) for both AMT cruises, at the CSIRO National Collections and Marine Infrastructure Data Trawler (https://www.cmar.csiro.au/data/trawler/survey_details.cfm?survey=IN2016_V01) for IND- 2017 and HEOBI, at the CLIVAR and Carbon Hydrographic Data Office (https://cchdo.ucsd.edu/) for GO-SHIP P15S, P16N and P16S, at the Australian Antarctic Division Datacenter (https://data.aad.gov.au/aadc/voyages/ https://doi.org/10.6075/J0CCHLY9) for the I08S and I09N cruises, at the MGDS (Marine Geoscience Data System: https://www.marine-geo.org) for POTATOE, and finally at (https://scope.soest.hawaii.edu/data/gradients/documents/) for both SCOPE-Gradients cruises. Finally, we used satellite data to estimate the euphotic zone depth where photosynthetic available radiation (PAR) is 1% of its surface value^27,28^. We approximated the euphotic zone depth using the light attenuation at 490 nm (Kd 490) product and the relationship Z eu (1%) (euphotic zone depth) = 4.6/Kd 490 (diffuse attenuation coefficient for downward irradiance). We also used the script Longhurst-Province-Finder (https://github.com/thechisholmlab/Longhurst-Province-Finder) to assign each sample to the Longhurst Province in which it was sampled; helping subset data and investigate specific regions of the ocean.

### DNA Extraction

For AMT cruises, DNA was isolated and purified using the AllPrep DNA/RNA Mini kit (QIAGEN) with modifications to be compatible with RNAlater® and to disrupt cell membranes^29^. Briefly, the filter was removed from the Sterivex housing and immersed in RLT buffer that had been amended with 10 µl 1 N NaOH per 1 ml buffer, followed by a 2 min agitation in a Mini-Beadbeater-96 (Biospec) with 0.1 and 0.5 mm sterile glass beads (BioSpec). DNA was isolated and purified from the FRAM and MOSAiC samples using the PowerWater kit (QIAGEN), while DNA from bioGEOTRACES was isolated and purified using a phenol/chloroform-based method^30^, and DNA from the SCOPE-Gradients cruises was isolated and purified following^31^. DNA from IND- 2017, K-AXIS, HEOBI, and P15S was isolated and purified, using a modified phenol:chloroform:isoamyl based extraction protocol of the DNeasy PowerWater Sterivex Kit^32–34^. For I09N, I08S, P16N, P16S, and POTATOE, the DNA was isolated and purified using the method described in^35^, with modifications detailed in^36,37^. An additional bead-beating step was added using 0.1 mm glass beads (Biospec) for two minutes at maximum speed on a VWR Analog Vortex Mixer. DNA was precipitated in low-EDTA buffer to protect from degradation and quantified using PicoGreen dsDNA assay (Thermo Fisher Scientific). A detailed extraction protocol is available at https://osf.io/k4c5u.

### Sequencing methods and pipeline

Relative abundance information was generated by PCR amplification of extracted DNA using the 515Y (5ʹ-GTGYCAGCMGCCGCGGTAA), 926R (5ʹ-CCGYCAATTYMTTTRAGTTT) universal rRNA gene primer pair^20^ which amplifies both 16S and 18S rRNA simultaneously. All DNA samples, were amplified using the protocol available at (https://www.protocols.io/view/fuhrman-lab-515f-926r-16s-and-18s-rrna-gene-sequen-vb7e2rn), with the exception of FRAM, for which raw fastq files were supplied. Briefly, PCR master mix consisted of 12 µl of PCR water (VWR), 10 µl of 5’ Master Mix (0.5 U Taq, 45 mM KCl, 2.5 mM Mg^2+^, 200 µM each dNTP), 1.5 µl of 1:1, 515F:926R barcoded primer mix (0.3 mM each primer), and 1 µl of DNA per reaction for a final volume of 25 µl. We used 5’ master mix (Quantabio) for I8S, I9N, P16N, P16S, AMT-19, AMT-20, POTATOE, GA02, GA03, GA10, GP13, SCOPE-Gradients 2, and SCOPE-Gradients 3 samples, GoTaq master mix (Promega) for P15S, K-AXIS, IND-2017, and HEOBI samples, and U Phusion Polymerase (Thermo Fisher Scientific) was used by collaborators for FRAM and MOSAiC samples. PCR was performed under the following conditions: initial denaturing at 95°C for 120 seconds, followed by 30 cycles of 95°C for 45 seconds, 50°C for 45 seconds, and 68°C for 90 seconds, and then a final elongation step at 68°C for 300 seconds. PCR products were stored at 4°C before being cleaned using the Agencourt AMPure XP PCR purification protocol. Finally, DNA concentration was quantified using the Pico-Green dsDNA Quant-iT Assay Kit. We used barcoded rRNA gene primers that had linkers, so they made sequencing libraries directly. Pooled sequences for each run were cleaned and concentrated with SPRIselect (Beckman Coulter) beads. The cleaned pool was quantified using the Qubit dsDNA HS Assay Kit. Finally, the 16S and 18S rRNA amplicon concentrations in the pool were determined with a Bioanalyzer Chip: High Sensitivity DNA Kit, and their ratio used for later data correction.

Most sequencing was done using HiSeq Rapid Run technology (2 x 250 bp), except for FRAM and MOSAiC which were sequenced using MiSeq technology (2 x 300 bp). Sequences were demultiplexed manually (https://github.com/jcmcnch/demux-notes) and denoised to amplicon sequence variants (ASVs) with a custom analysis pipeline based on QIIME2 and DADA2^21,22^ which can be found on GitHub (https://github.com/jcmcnch/eASV-pipeline-for-515Y-926R). The main difference between this pipeline and standard workflows is that it contains an initial 16S/18S rRNA splitting step, which is accomplished using bbsplit against curated 16S and 18S rRNA databases derived from SILVA version 138.1 and PR2 version 4.14.0. This results in two subsets of data (16S and 18S rRNA) that are then denoised separately and later merged. Note that standard pipelines require a merging of forward and reverse reads where they overlap, which for these rRNA gene primers would lead to the removal of all 18S rRNA sequences because their forward and reverse 18S rRNA reads do not overlap. While the standard method (with overlap merging) was applied to the 16S sequences, the 18S rRNA sequences were concatenated. We used a universal trim length of 220 bp for forward and 180 bp for reverse reads before concatenation so that ASVs could be directly compared to each other across cruises. After denoising, sequencing depth was calculated, which ranged from 1 – 1, 250, 359, and we filtered any samples with a sequencing depth below 5000 out of the dataset, as they were considered too small to be statistically useful. On average, there were 180, 528 sequences per sample before implementing the two corrections described below.

Two corrections were made to both 16S and 18S rRNA ASV tables before they were merged. The first adjusts for a known bias against longer 18S rRNA sequences during Illumina sequencing^23^ and the second adjusts for random variations in sample quality using the DADA2 output statistics. Briefly, for the first adjustment, bioanalyzer traces were taken for all samples except FRAM and MOSAiC samples to determine the fraction of 16S rRNA or 18S rRNA prior to sequencing, which was calculated by:

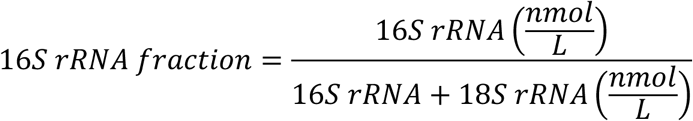

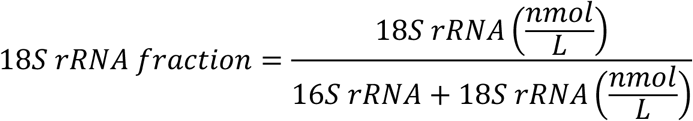

This was considered the “expected” outcome from the sequencing run. The same fractions were then calculated after the bbsplit step (which sorts 16S rRNA and 18S rRNA sequences into separate bins) in the analysis pipeline (Figure 2), using the number of 16S and 18S rRNA sequences measured by the sequencer. These ratios were calculated by:

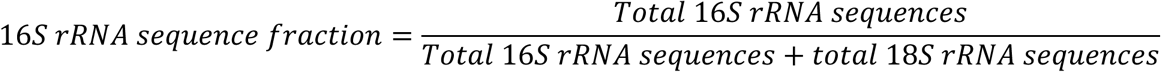

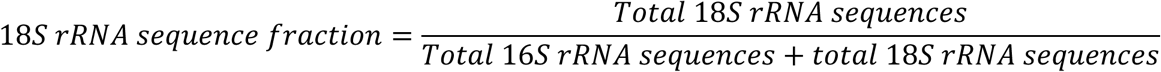

**Figure 2:**
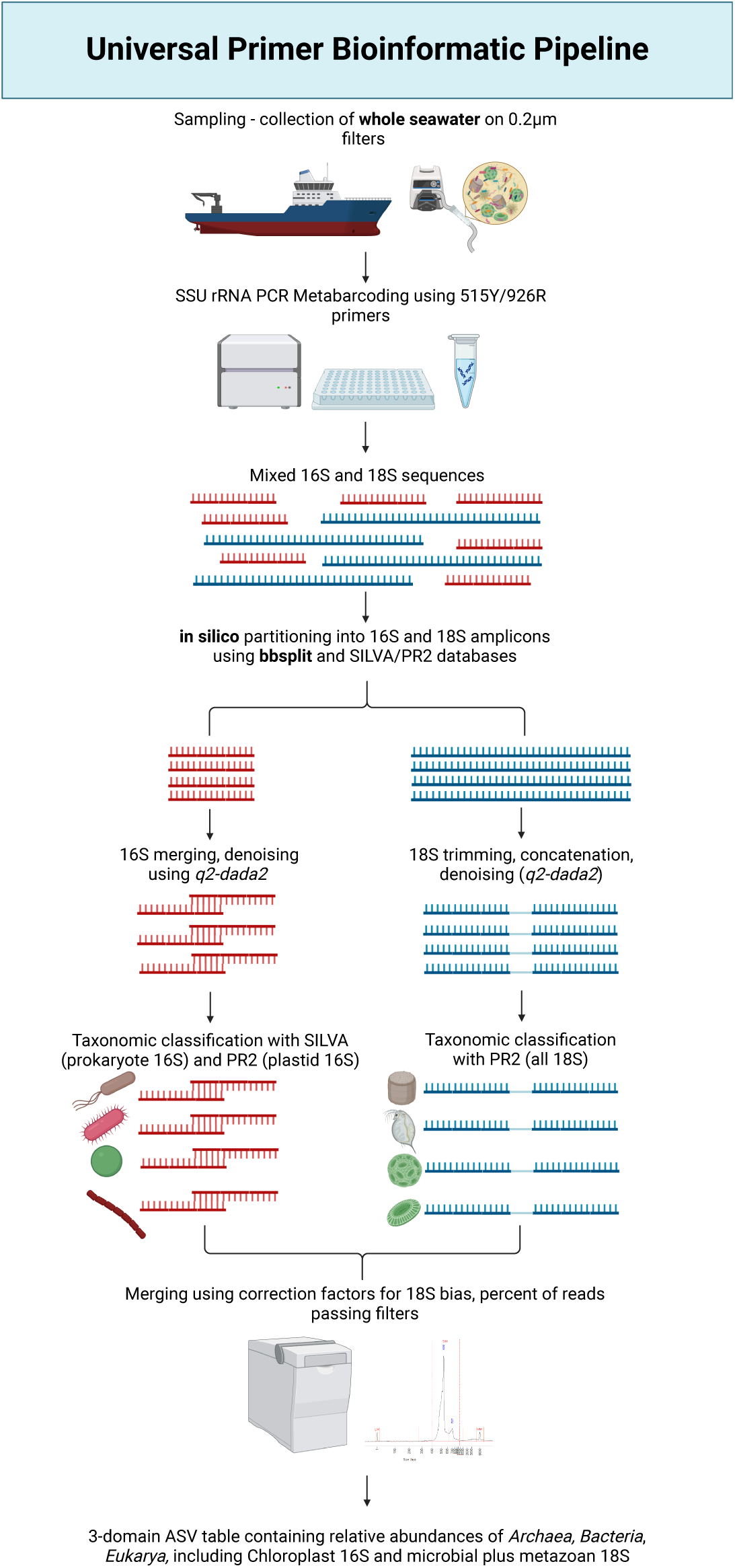
Bioinformatic pipeline used to analyze mixed 16S and 18S rRNA amplicons generated from three-domain universal SSU rRNA primer set 515Y/926R from unfractionated seawater collected in the GRUMP collaboration. The unique features of this pipeline include an initial database-dependent *in silico* splitting step after PCR amplification and sequencing, parallel 16S rRNA and 18S rRNA denoising pipelines, and a correction-based merging step − allowing for the recovery of both 16S rRNA and 18S rRNA information in a single ASV table. Note that separate analysis pipelines are required due to the non-overlapping nature of forward and reverse 18S rRNA reads which would otherwise cause them to be discarded in bioinformatic pipelines that are dependent on read merging (as is common for most rRNA-based amplicon pipelines). The pipeline is based primarily in the QIIME2 framework and has been implemented and tested with Ubuntu-based Linux distributions.

A correction factor for each sequence type was calculated as:

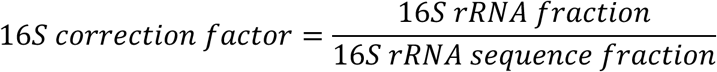

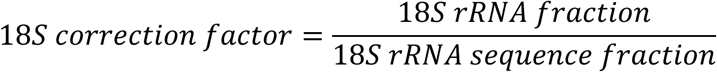

The second adjustment for random variations in sample quality uses the DADA2 output statistics. The total number of non-chimeric reads were divided by the initial number of input reads, which is the ratio of reads that passed the DADA2 chimera detecting software. We then corrected our output read number for each sample by the following equation:

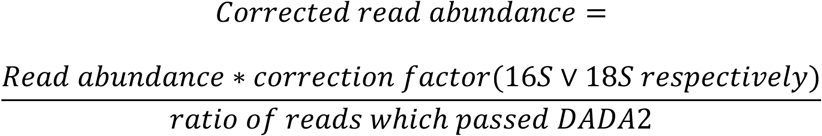

All of these calculations were made using the script available at https://github.com/Nwilliams96/Project-2-Universal-Primer-Pipeline. FRAM and MOSAiC data does not have bioanalyzer traces; we used the same correction factor used for the Southern Ocean samples, as we hypothesize, they would have a similar level of eukaryotic bias. The final dataset contains merged abundance information from both 16S rRNA ASVs and 18S rRNA ASVs that can be directly compared to one another. Correction factors and calculations are available in Supplementary Table 2.

In the final dataset, metabarcode information was derived from bacteria and archaea (16S rRNA), protists (photosynthetic based on plastid 16S rRNA and non-photosynthetic based on nuclear 18S rRNA), as well as larger planktonic organisms such as arthropods, salps, and gelatinous zooplankton (nuclear 18S rRNA), which are derived from tissue fragments, larvae, individuals, eggs, etc. that were captured on 0.2 µm filters. Note that both chloroplast 16S rRNA and nuclear 18S rRNA will represent most photosynthetic protists, with the exception of dinoflagellates as they do not all have recognizable or amplifiable chloroplast sequences (Lila Koumandou et al., 2004 and personal communication with Sanchez-Garcia et al. (Sanchez-Garcia et al., 2022)). The taxonomy levels are joined from three databases: PR2^40^, SILVA^41^, and ProPortal^42^. The levels are described in Table 1.

**Table 1.**
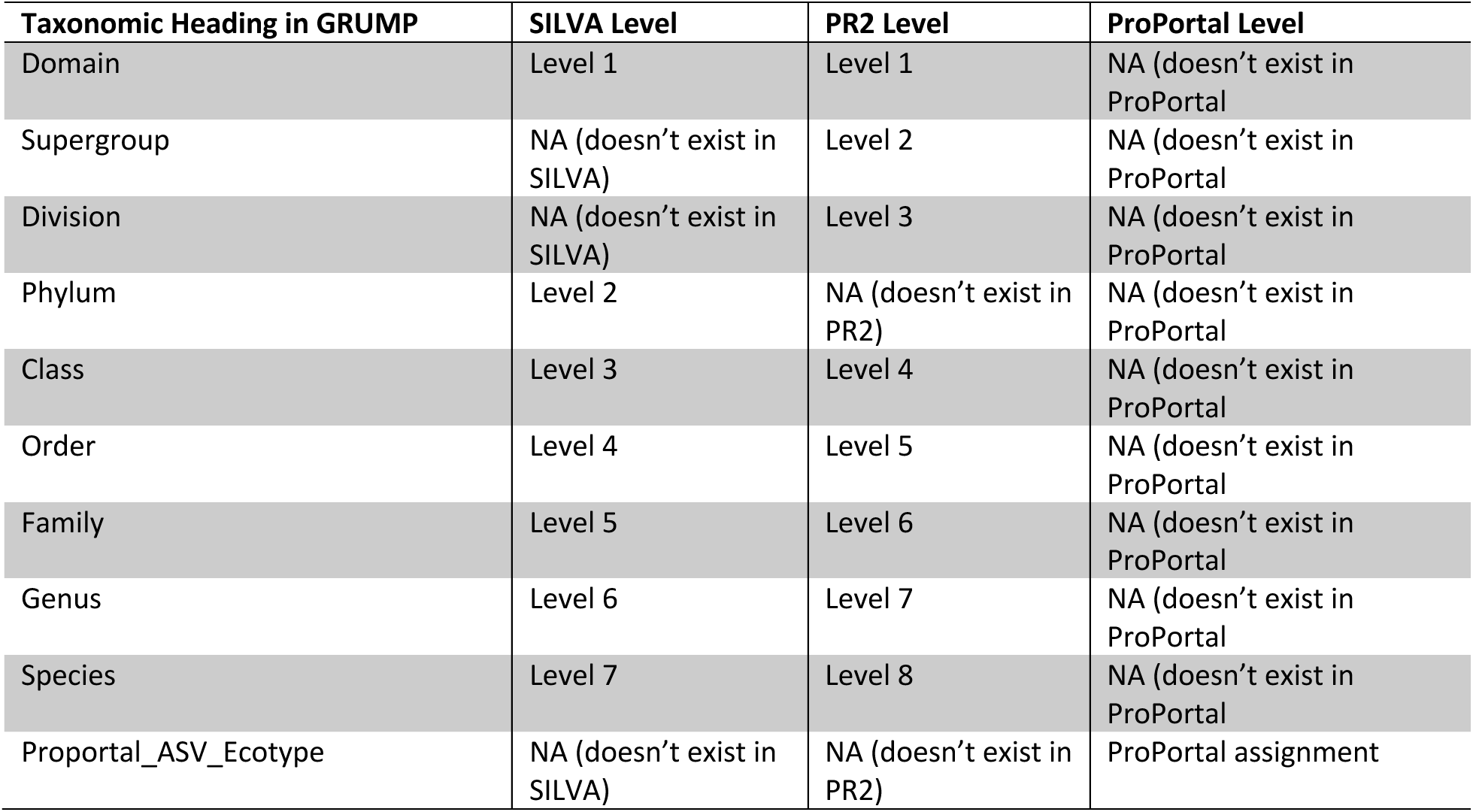
Taxonomic headings in the GRUMP dataset, with their corresponding database levels. This was necessary as not all levels of taxonomy are aligned across databases.

| **Taxonomic Heading in GRUMP** | **SILVA Level** | **PR2 Level** | **ProPortal Level** |
| --- | --- | --- | --- |
| Domain | Level 1 | Level 1 | NA (doesn't exist in ProPortal) |
| Supergroup | NA (doesn't exist in SILVA) | Level 2 | NA (doesn't exist in ProPortal) |
| Division | NA (doesn't exist in SILVA) | Level 3 | NA (doesn't exist in ProPortal) |
| Phylum | Level 2 | NA (doesn't exist in PR2) | NA (doesn't exist in ProPortal) |
| Class | Level 3 | Level 4 | NA (doesn't exist in ProPortal) |
| Order | Level 4 | Level 5 | NA (doesn't exist in ProPortal) |
| Family | Level 5 | Level 6 | NA (doesn't exist in ProPortal) |
| Genus | Level 6 | Level 7 | NA (doesn't exist in ProPortal) |
| Species | Level 7 | Level 8 | NA (doesn't exist in ProPortal) |
| Proportal_ASV_Ecotype | NA (doesn't exist in SILVA) | NA (doesn't exist in PR2) | ProPortal assignment |

The forward and reverse sequences were submitted to the NCBI database under accession numbers GA02/GA10: PRJNA1194189, GA03: PRJNA1194192, GP13: PRJNA1195113, Scope-Gradients 2: PRJNA1195115, Scope-Gradients 3: PRJNA1196422, P16N: PRJNA1196483, P16S: PRJNA1196490, P15S: PRJNA1196498, HEOBI: PRJNA1196504, IND-2017: PRJNA1196513, K-AXIS: PRJNA119651, POTATOE: PRJNA1198072, I08S: PRJNA1198088, I09N: PRJNA1198607, MOSAiC: PRJNA1198992, and FRAM: PRJNA1199044. The scripts necessary to exactly reproduce the analysis are available at https://github.com/jcmcnch/Global-rRNA-Univeral-Metabarcoding-of-Plankton, with modifications to the merge script found at https://github.com/Nwilliams96/Project-2-Universal-Primer-Pipeline-Modifications.

### 2.4 Statistical Analysis

All statistical analyses and visualizations were performed in R v4.3.2^43^ using vegan^44^, tidyverse^45^, ggpubr^46^ and patchwork^47^ packages. All packages were used with their default settings. Scripts used to arrange data, perform statistics, and create figures can be found at (https://github.com/Nwilliams96/Project-1-GRUMP-data-release).

## Data Records

### Dataset Overview

The Global rRNA Universal Metabarcoding Plankton database (GRUMP) is currently at version 1.3.3. GRUMP version 1.3.3 is a collection of 1194 metabarcoding community profiles from 19 different oceanographic samplings from all major ocean basins, covering a wide depth, oceanographic, and seasonal range (Table 2, Figure 1 A and B). The dataset covers 29 Longhurst provinces, ranging from 0 - 6015 m in depth, and dates between 2003 and 2020.

**Table 2.** Summary of GRUMP. Note cruises spanning the equator covered opposite seasons depending on location.

| **Cruise** | **Ocean** | **Depth**<br>**Range (m)** | **Time Frame** | **Season(s)** | **Method of Sampling** | **Volume**<br>**filtered** |
| --- | --- | --- | --- | --- | --- | --- |
| **I8S** | Indian | 6 – 1010 | 18/2/07 – 27/2/07 | Winter & Summer | Niskin | 2 L |
| **I9N** | Indian/Southern | 14 – 1450 | 1/3/07 – 27/4/07 | Autumn | Niskin | 2 L |
| **POTATOE** | Pacific | Surface Only | 22/08/03 – 11/10/03 | Spring & Summer | Bucket | 20 L |
| **SCOPE-Gradients 2** | Pacific | 12 – 120 | 26/5/17 – 13/6/17 | Spring & Summer | Niskin & underway | 0.7-4 L |
| **SCOPE-Gradients 3** | Pacific | Surface – 125 | 11/4/19 – 28/4/19 | Spring | Niskin & underway | 0.7-4 L |
| **P16N** | Pacific | 20 – 1003 | 15/2/06 – 26/3/06 | Winter | Niskin | 2 L |
| **P16S** | Pacific/Southern | 2 – 500 | 11/1/05 – 9/2/05 | Summer | Niskin | 2 L |
| **P15S** | Pacific/Southern | 4.8 – 6015 | 4/5/16 – 24/6/16 | Autumn & Winter | Niskin | 2 L |
| **bioGEOTRACES**<br>**GA02** | Atlantic | 8 – 202 | 11/05/10 – 27/3/11 | Summer | Niskin | 100 ml |
| **bioGEOTRACES**<br>**GA03** | Atlantic | 2 – 1055 | 24/10/10 – 10/12/11 | Winter | Niskin | 100 ml |
| **bioGEOTRACES**<br>**GA10** | Atlantic | 6 – 121 | 19/10/10 – 15/11/10 | Autumn & Winter | Niskin | 100 ml |
| **bioGEOTRACES**<br>**GP13** | Atlantic | 14 – 5600 | 14/5/11 – 21/6/11 | Winter | Niskin | 100 ml |
| **AMT-19** | Atlantic | Surface Only | 20/10/09 – 25/11/09 | Autumn & Spring | Niskin | 5-20 L |
| **AMT-20** | Atlantic | Surface Only | 19/10/10 – 17/11/10 | Autumn & Spring | Niskin | 5-20 L |
| **IND2017** | Southern | 5 | 16/1/17 – 7/2/17 | Summer | Niskin | 2 L |
| **K-AXIS** | Southern | Surface Only | 11/1/16 – 22/1/16 | Summer | Niskin | 2 L |
| **HEOBI** | Southern | 8 – 2297 | 12/1/16 – 31/1/16 | Summer | Niskin | 2 L |
| **FRAM** | Arctic | 30-80 | 8/1/16 – 6/11/17 | Year-round | Remote Access<br>Sampler | 700 ml |
| **MOSAic** | Arctic | 736 – 5111 | 11/11/19 – 9/7/20 | Winter, Spring<br>and Summer | Niskin | 1-4 L |

Profiles were obtained through collaborations with researchers from around the world who performed extensive basin-scale oceanographic surveys. Whole seawater (> 0.2 µm) DNA extracts were amplified with the 515Y/926R three-domain universal metabarcoding primer set with conserved binding sites in both prokaryotic and eukaryotic small subunit (SSU) rRNA, which also includes chloroplast organellar 16S rRNA^20^. This primer set produces comprehensive planktonic community profiles from **<u>A</u>**rchaea to **<u>Z</u>**ooplankton (and beyond) at high taxonomic resolution. This is achieved using modern denoising algorithms that can differentiate amplicons into single-nucleotide variants^22^. Due to the unfractionated nature of the DNA extracts and the extensive organismal coverage achieved with the 515Y/926R rRNA gene primers, GRUMP integrates insights across trophic levels from many individual sampling efforts.

GRUMP includes high-quality environmental covariates to contextualize metabarcoding community profiles, easily accessible through the Simons Collaborative Marine Atlas Project (CMAP) database^48^ for simple downloading and analysis by non-experts. The core set of environmental covariates include temperature, salinity, and oxygen concentration, and for several datasets, silicate (SiO4^2-^), nitrite (NO2^-^), nitrate (NO3^-^), ammonium (NH4^+^), phosphate (PO4^3-^), dissolved organic carbon (DOC), total organic carbon (TOC), photosynthetically active radiation (PAR), and chlorophyll-*a*. While analysis of these variables is outside the scope of this data release, the full oceanographic dataset presents a key advantage for biogeographic studies by interpreting microbial distribution in the environmental context.

Pioneering studies such as Malaspina^14^ and TARA^13,49^ extensively sampled from the Atlantic, Pacific, Indian, Arctic, and Southern Oceans, as well as the Mediterranean and Red Seas.

GRUMP is distinguished by a uniquely large coverage of samples within the Southern and Pacific Oceans, with 251 and 510 samples respectively (Figure 1) and incorporation of samples from oceanic regions where there is a paucity of data. Furthermore, much of the GRUMP collaborative samples traverse long meridional transects and biological provinces (e.g. Alaska to Antarctica). Ultimately, we envision GRUMP as an evolving dataset to which samples will be added, facilitating comparisons between transects over time. TARA and Malaspina also used size-fractionated filtration to separately capture viruses (0.02 – 0.22 µm), free-living bacteria and archaea (0.2 – 1.6 µm), small eukaryotes and aggregated prokaryotes (0.8 – 5 µm), and large eukaryotes (5 – 20 µm and 20-2000 µm). Fractionating samples is a powerful approach that enhances the depth coverage of targeted organisms, providing an analysis that can improve sequencing efficiency (for organisms of interest) by reducing the complexity of the sequencing data and reducing unwanted organisms from that sample. Notably, sequencing results from those size fractions were compositional, meaning taxa were reported as a proportion of that size fraction alone − a major impediment to comparing between size fractions. Within GRUMP, we sequenced unfractionated samples using universal rRNA gene primers, which capture the entire microbial community so that results from bacteria, archaea and eukaryotes can be directly compared to one another. This will provide new broad insights to the global ocean’s microbiology, as significant contextual quantitative information is lost when considering size fractions independently. One area that unfractionated data are particularly important is the study of the many interactions among the microbial community, within and across trophic levels. For example, interactions between prokaryotic heterotrophs and eukaryotic phytoplankton ^50^ are thought to impact major elemental fluxes and the biological carbon pump^51^.

### Unique Technical Aspects

Due to the nature of the collaborative work, DNA extractions were carried out with diverse extraction procedures on whole seawater samples. For the samples extracted at USC (I08S, I09N, P16N, P16S, and POTATOE), the extraction procedure has been reported to have nearly 100% recovery of environmental microbial nucleic acids^35^, and we included an additional bead-beating step to lyse recalcitrant algal cells and metazoan tissue.

With the exception of FRAM and MOSAiC data, all extracts were amplified using a 1-step PCR protocol (https://www.protocols.io/view/fuhrman-lab-515f-926r-16s-and-18s-rrna-gene-sequen-j8nlkpd1g5r7/v2) where PCR rRNA gene primers are ordered as pre-ligated constructs that contain Illumina adapters, sequencing rRNA gene primers, and barcodes (https://osf.io/57dpa/). This approach requires less pipetting and fewer consumables (including PCR reagents) versus 2-step protocols and has the key advantage of minimizing potential cross-contamination since barcodes are already integrated within sequencing libraries after PCR amplification. Users intending to apply this protocol to their samples should note two caveats: firstly, the libraries produced with this method require approximately 3 times the normal loading concentration versus 2-step protocols, likely due to inefficient clustering on Illumina flow cells for unknown reasons. Recent experience suggests that optimal clustering on first-generation MiSeq platforms (i.e. bridge amplification, not more recent patterned flow-cell technology) can be obtained with the maximum manufacturer-recommended loading concentration of 20 pM for amplicon pools with 10% phiX. Secondly, the 1-step primer constructs used in this protocol (with a combinatorial barcoding scheme, i.e. each barcoded forward primer can be matched with multiple barcoded reverse rRNA gene primers) are only appropriate for bridge amplification sequencing (i.e. MiSeq, Aviti, and the now obsolete HiSeq), not patterned flow cells (e.g. NovaSeq and other more recent Illumina technologies). With the latter, unique dual indexes (UDIs) are required to avoid the impact of higher “index hopping” inherent to patterned flow cell technology.

All amplicon sequences were analyzed with a custom bioinformatics pipeline that is designed for 3-domain metabarcoding data (Figure 2). This pipeline’s default settings are for the 515Y/926R primer set because with these rRNA gene primers, prokaryotes have overlapping forward and reverse reads, but the longer eukaryote amplicons require concatenation to avoid discarding all eukaryote sequences lacking overlap (merging overlapping forward and reverse reads via their overlap is standard in virtually all amplicon pipelines). The unique feature of this 3-domain analysis workflow is the ability to split mixed 16S rRNA and 18S rRNA amplicon sequences into two separate pools, which are analyzed separately (16S rRNA forward and reverse merged, 18S rRNA forward and reverse concatenated). The output of these two analyses are then merged into a single 3-domain table using correction factors described above. The pipeline also automates routine analysis from demultiplexed fastq files, including installation and database setup, primer trimming, 16S/18S rRNA splitting, denoising, taxonomic annotation, and data import/export to the QIIME2 framework.

### GRUMP Contextual Data: Ecologically Relevant Taxonomic Annotations & Groupings

In addition to the core set of environmental covariates discussed above, we have also provided grouping of sequences into ecologically relevant plankton groups, which have been curated based the most abundant and highly diverse groups within the dataset. These ecologically relevant plankton groups do not adhere to usual taxonomic levels, and so we believe they are a particularly useful and practical way to subset the data. For example, if one was interested in comparing the abundance of a eukaryotic phytoplankton to the abundance of *Prochlorococcus* or *Synechococcus* within any given sample, the data can directly be subset using the “Eco_relevant_plank_groups” column within the dataset. For ecologically relevant plankton groups see Figure 3.

**Figure 3.**
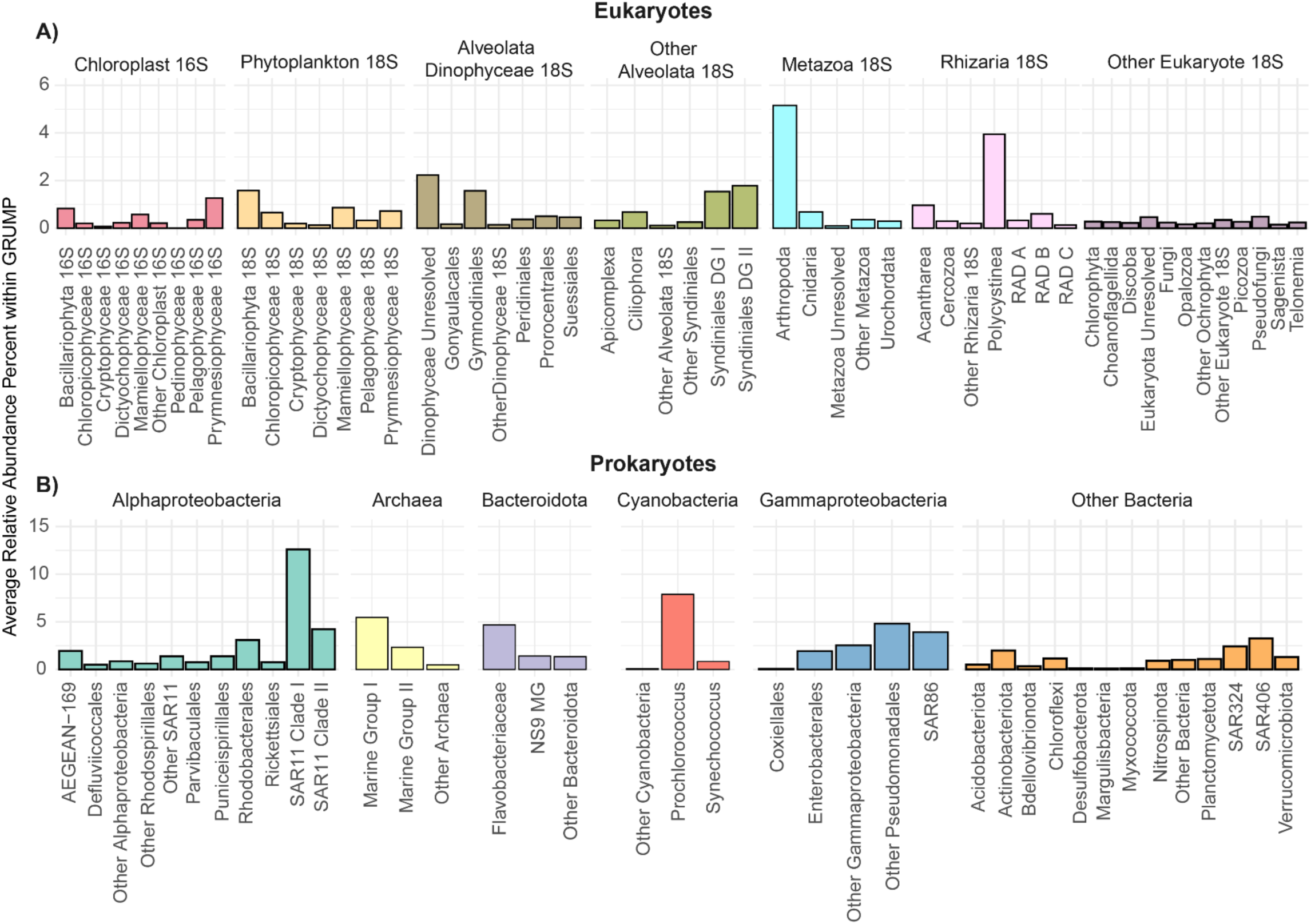
The average relative abundance from major marine plankton groups across all GRUMP samples. **A)** Eukaryotes and **B)** Prokaryotes.

We extracted all cyanobacteria ASV sequences and re-annotated them using the highly- resolving cyanobacteria database ProPortal^42^. ProPortal is currently consists of 1110 *Prochlorococcus* and 160 *Synechococcus* isolate and single cell genomes, which allowed us to resolve 69 ASVs into *Prochlorococcus* and 10 ASVs into *Synechococcus* ecotypes. This is a higher resolution than has traditionally been thought possible with 16S rRNA amplicons.

### Overall Taxonomic Composition of GRUMP Dataset

We observed 136,603 unique ASVs, and amongst these unique ASVs, 359,030,338 reads, after DADA2 processing. Of these raw reads, 335,265,166 were prokaryotic and 23,765,172 were eukaryotic (including chloroplast 16S). To interpret the results quantitatively, it should be considered that Illumina sequencers exhibit a bias toward shorter fragments. As a result, the 18S rRNA amplicons (which can be hundreds of bases longer than the 16S rRNA amplicons generated from these rRNA gene primers) are under-represented. We correct for this bias by comparing the relative abundances of 16S and 18S rRNA amplicons in the original sequencing libraries (measured by Bioanalyzer) with the final proportion of 16S and 18S rRNA sequences after DADA2 processing. We apply a correction factor to each sequence type, calculated as the proportion of the sequence types reads in the bioanalyzer output divided by the proportion in the final sequences. This empirical correction factor ranged from 1.90 to 9.42 (18S rRNA) and 0.75 to 0.90 (16S rRNA). Once adjusted for this bias against eukaryotes, the total corrected number of reads was 256,299,695 with 165,518,629 prokaryotic reads (17,443,053 archaea, 165,518,629 bacteria) and 72,455,193 eukaryotic reads, a global ocean ratio of roughly 2:1. Data users should note that this correction results in non-integer values for raw read counts in the corrected output tables.

Among the 136,603 ASVs classified in GRUMP, we observed 9 supergroups, 34 divisions, 66 phyla, 307 classes, 671 orders, 1222 families, 2256 genera, and 2030 species. Many of the species were represented by hundreds of ASVs. Further, use of the ProPortal database^42^ identified an additional 27 *Prochlorococcus* and *Synechococcus* ecotypes. Currently within the PR2 database^40^, there are 9 super groups, 33 divisions (of which we observed all), and 48,202 species, of which we only observed 1162. Comparatively the SILVA database^41^ has 94 prokaryotic phyla, of which we observed 66, and over a million prokaryotic species of which we observed 871, keeping in mind many species were represented by hundreds of ASVs, and 59,383 ASV assignments were truncated at a coarser level than species.

To gain a more resolved snapshot of the GRUMP dataset, we allocated the ASVs from both fractions into ecologically relevant categories based on prior marine microbial diversity studies (Figure 3). We divided the prokaryotic ASVs into six categories at the same level, Alphaproteobacteria, Archaea, Bacteriodota, Cyanobacteria, Gammaproteobacteria, and an “Other Bacteria” category. This allows users to compare these six groups to one another, rather than at the rigid levels provided by SILVA and PR2. The most abundant ecologically relevant group of archaea was Marine Group I, with an average relative abundance of 5.45%, more than double the average relative abundance of Marine Group II. Alphaproteobacteria had both the most unique ASVs (27,002) and the highest number of identified reads, resulting in an average relative abundance of 28%. SAR11 Clade I was the most abundant clade of Alphaproteobacteria, had an average relative abundance of 12.6%. The cyanobacterium *Prochlorococcus* had a total of 607 unique ASVs and average relative abundance of 7.8% across GRUMP. One significant advantage of this dataset is that due to the long amplicon covering two hypervariable regions, the majority of these ASVs could be annotated at fairly high phylogenetic resolution with the marine cyanobacteria genomic database ProPortal^42^, which allows for investigation of these organisms at the ecotype level, paving the way for researchers to ask specific questions about the biogeography and niches of the *Prochlorococcus* ecotypes at a global scale.

Eukaryotic phytoplankton were divided into two groups: one derived from 16S rRNA chloroplast sequences, and one derived from 18S rRNA sequences. In general, there were over 2.5 times more chloroplast 16S rRNA reads and 7 times more individual chloroplast 16S rRNA ASVs than 18S rRNA phytoplankton ASVs. Multiple chloroplast variants relative to a single 18S rRNA ASV has been previously reported for the South China Sea^52^ and at the San Pedro Ocean Time Series (SPOT)^53^. In our dataset, a prominent example of this is from the Prymnesiophytes for which we obtained 3,978 ASVs from the chloroplast 16S and 607 from the 18S rRNA. This could explain why this group of phytoplankton is much more abundant across the dataset when looking at the chloroplast 16S rRNA, with an average relative abundance of 1.2%, compared to an average relative abundance of 0.7% from the 18S rRNA sequences. So rather than being a discrepancy, this point highlights a major strength of using these universal rRNA gene primers, which detect both sequence types, and enable the user to select the appropriate data that will best answer their ecological question.

The most abundant ecologically relevant plankton group amongst the larger eukaryotes was Arthropoda, which, whilst having an average relative abundance of 5.15% across all GRUMP samples, had many fewer ASVs than most other groups (665). Amongst the Arthropoda, 95% of the reads and 90% of the unique ASVs were from Maxillopoda. This is also the case in SPOT data, where it has been argued that most of these sequences were likely from juveniles, eggs or organismal fragments, given that these SPOT data were fractionated between 1 – 80 µm and that these organisms are generally larger than 80 µm as adults^10^. However, given GRUMP is unfractionated and not pre-filtered, some of these sequences could be from adult organisms. Of note, there are a substantial portion of the Arthropoda that are identified down to the species level, such as *Cyclopina gracilis*, a marine copepod, highlighting the range at which this dataset can be of use when analyzing a variety of organisms. Notably, different species of marine copepods can have a wide range of 18S rDNA copy number (> 1,000) and thus, the relative abundances of these organisms require cautious interpretation^54^.

### Spatiotemporal Trends of Community Composition Across GRUMP

Of the total 256,299,695 reads, 8.8% were from the Arctic Ocean, 17.3% from the Atlantic Ocean, 9% from the Indian Ocean, 39.5% from the Pacific Ocean, and 25.5% from the Southern Ocean (Figure 4 A). By depth, 87.3% of the reads were from surface to 200 m, 10.9% from 200 – 1000 m, and 1.9% from below 1000 m (Figure 4 B). Within the 136,603 total ASVs, 50% (68,996) were unique to a single sample, whereas no ASVs was detected in every sample. At the ocean basin scale, 4.1%, 12.4%, 10.1%, 27.8%, and 12.9% of the ASVs were unique to the Arctic, Atlantic, Indian, Pacific, and Southern Ocean, respectively, and 0.77% (1046) were found within all five oceans. The most abundant ASVs observed in all ocean basins were those assigned to three HLII *Prochlorococcus*, SAR11 Clade I, SAR11 Clade II, and Maxillopoda.

**Figure 4.**
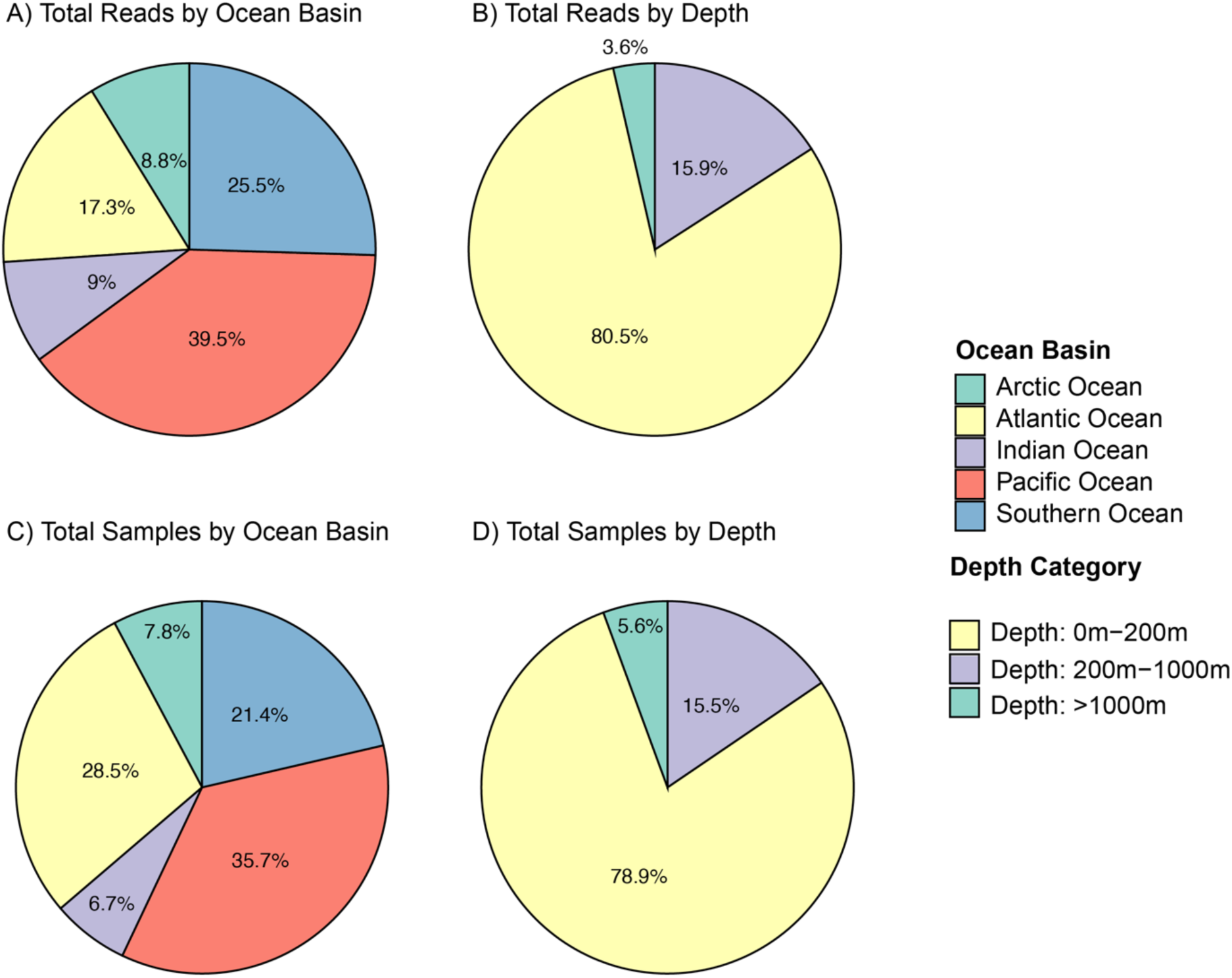
Percent of total reads derived from **A)** each ocean basin or **B)** various depths. Total reads in GRUMP: 256,299,695.

While it is beyond the scope of this report to go into detail about the distributions of most taxa at fine phylogenetic resolution, it is illuminating to divide the results into five major plankton groups including the archaea, bacteria (minus cyanobacteria), cyanobacteria, chloroplast 16S (eukaryotic phytoplankton) and eukaryotic 18S. In the top 50 m of the ocean, the most abundant of these five major plankton groups was bacteria (minus cyanobacteria) (Figure 5 B), contributing a mean relative abundance of 62% in the Pacific Ocean, 60% in the Indian Ocean, and 72% in the Atlantic Ocean. The mean relative abundance of bacteria in the polar oceans was much lower with an average relative abundance of 36% in the Arctic and 54% in the Southern Ocean (Figure 5 A, C). Of the phytoplankton, the eukaryotic phytoplankton (via chloroplasts) were observed at a similar relative abundance throughout the Pacific, Atlantic, and Indian oceans, whereby the mean relative abundances were 4%, 3%, and 2% respectively, while in the polar oceans the average relative abundance was 5% in the Arctic and, 9% in the Southern Ocean, which is more than double what was observed in the Pacific, and 4 times what was observed in the Indian (Figure 5 A). The cyanobacteria were much more abundant than their eukaryotic phytoplankton counterparts in the top 50 m of the ocean whereby the average relative abundance in the Pacific Ocean was 14%, the Indian Ocean 24%, and the Atlantic Ocean 9%, but were only rarely detected in the polar oceans, and then generally at a very low abundance (Southern Ocean mean = 1%, Arctic Ocean mean = 0.23%). In the top 50 m, the archaea were the least abundant of these five groups, contributing between a mean relative abundance of 1% in the Indian Ocean, to a mean of 2% in the Southern Ocean (Figure 5). For depth figures, please see Supplementary Figures 2 – 5.

**Figure 5.**
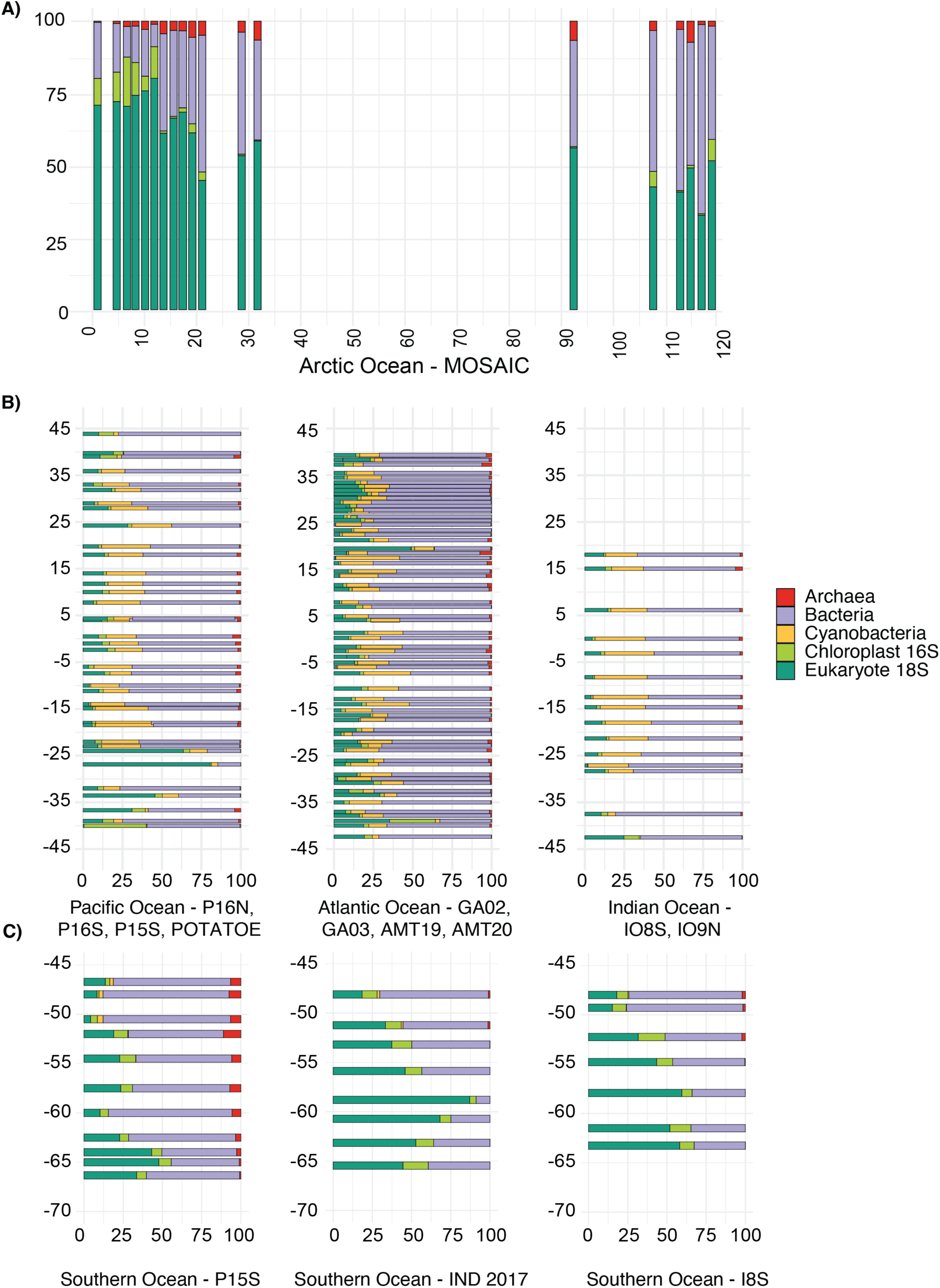
Relative abundance of marine microbes in the top 25 m of the ocean, partitioned between archaea, bacteria, cyanobacteria, chloroplast 16S rRNA, and eukaryotic 18S. A) Relative abundance % (y axis) across longitude in decimal degrees (x axis) from MOSAiC within the Arctic Ocean. B) Relative abundance % (x axis) vs latitude in decimal degrees (y axis) from within the Pacific Ocean, the Atlantic Ocean, and Indian Ocean C) relative abundance (x axis) vs latitude in decimal degrees (y axis) from within the Southern Ocean.

### Potential Uses for the GRUMP Dataset

A main advantage of SSU rRNA community profiling (compared to shotgun metagenomes) is high coverage, whereby far more organisms are characterized and identified with much less sequencing than required by metagenomics. Virtually all the rRNA gene sequences are phylogenetically informative in a consistent framework. With metagenomes, only the fraction of sequences that map to references or MAGs, or whose short read match such references closely enough to identify, are useful for community composition analyses, and even then, sometimes at relatively coarse phylogenetic resolution^15^. In addition, with universal amplicons there is no assembly or other biases associated with MAG-based community compositions (note MAGs generally miss most SAR11 and *Prochlorococcus*, for example as they are hard to assemble due to their highly “microdiverse” genomes). Therefore, with SSU rRNA community profiling, the researcher gets broad, nearly comprehensive coverage, with the caveat that a small fraction of environmental sequences do not perfectly match the PCR rRNA gene primers and might be missed (ca 2% of prokaryotes and 10-15% of eukaryotes^24^). This means that this dataset is extremely powerful when answering ecological questions that require abundance information of ocean microbes, with detailed taxonomic resolution.

Beyond microbial ecological questions regarding biogeography and diversity, GRUMP can be used in a number of other ways. The contextual data provides the opportunity to evaluate how metabarcoding recovers spatiotemporal patterns in comparison to other techniques, such as flow cytometry ^55^, microscopy, pigments, etc. Regarding comparison of the metabarcoding data with pigments, the data might even be used to help calibrate and validate pigment data if collected at the same time. Finally, the data can also be used to compare and contrast patterns between real-world data and model outputs.

## Technical Validation

This rRNA gene primer pair (515F/926R) used in this study has been validated via both mock communities of marine bacteria, archaea^20^, and eukaryotes^23^; such accuracy is not observed with other commonly tested rRNA gene primers^23,24^. To ensure the integrity of each sequencing run, we included 16S, 18S, mixed and staggered mock communities as a positive control.

Our lab has also made comparisons with epipelagic metagenomes from around the world, which have shown that these primers perfectly match 96% of the corresponding genomic primer regions across all three domains^24^. Most importantly, this means that the relative abundance of amplicon sequencing variants (ASVs) from bacteria, archaea, and eukaryotes are able to be quantified together as a total community.

## Usage Notes

GRUMP is a collaborative effort that will continue to evolve. We have developed a multilayered data storage structure to accommodate future updates and changes. Simons CMAP will host processed relative abundance data alongside selected environmental covariates. CMAP is a user-friendly database that is accessible through a web interface and is suitable for users without special expertise in bioinformatics or oceanography. Direct download from web interface is possible. On CMAP, the dataset is in long format, without explicit zeroes. A GitHub page (https://github.com/jcmcnch/Global-rRNA-Univeral-Metabarcoding-of-Plankton) serves as a central repository for accessing code underlying the analysis, as well as raw metadata files and tables that may be useful for advanced users. We have also uploaded a file containing ASVs and their taxonomy to the Github as a BLAST database for users who may find this useful.

The OSF repository contains an archive of bioinformatic intermediates that may be useful for those wishing to analyze only portions of the dataset (e.g. 16S rRNA ASVs only, 18S rRNA ASVs only, or various subsets therein) as well as QIIME2 artifacts that can be used for reanalysis or other purposes. The raw, demultiplexed fastq files are available in the SRA. Note that limited metadata will be provided here. The GitHub page will always be the most up-to-date place to examine new metadata, followed by CMAP.

## Code Availability

All code is available on GitHub (https://github.com/jcmcnch/Global-rRNA-Univeral-Metabarcoding-of-Plankton and https://github.com/Nwilliams96/Project-2-Universal-Primer-Pipeline-Modifications).

## Acknowedgements

We would like to acknowledge the help of Laura Furtado (Fuhrman Lab Technician) for her assistance and support with lab work, sequencing, and organizing the postage of samples. We would also like to acknowledge the help of Bruce Yanpui Chan, as well as Yi-Chun Yeh for their support with DNA extractions, DNA amplification and sequencing.

We would like to acknowledge the help of many crew and technicians of various cruises on which data for this project were collected. For AMT, we would like to thank the captain and crew of the RRS James Cook, Stephanie Sargeant, and the marine technicians from UK National Marine Facilities for their assistance in collecting these samples. For FRAM and MOSAiC, we would like to thank the captain and crew of R/V Polarstern expeditions PS99.2 and PS107, as well as the FRAM / HAUSGARTEN team, for their exceptional support. We thank Jana Bäger, Theresa Hargesheimer, Rafael Stiens, Lili Hufnagel, Daniel Scholz, Normen Lochthofen, Janine Ludszuweit, Lennard Frommhold and Jonas Hagemann for RAS and mooring operation; Jakob Barz and Swantje Ziemann for DNA extraction and library preparation; Halina Tegetmeyer for amplicon sequencing; and Laura Wischnewski for nutrient analyses. For BioGEOTRACES we would like to thank the captain and crew of the R/V Pelagia, R/V Southern Surveyor, R/V Tangaroa, RRS James Cook, RRS Discovery, and R/V Knorr, and Kristin Bergauer, Heather A. Bouman, Thomas J. Browning, Daniele De Corte, Christel Hassler, Debbie Hulston, Jeremy E. Jacquot, Elizabeth W. Maas, Thomas Reinthaler, Eva Sintes, and Taichi Yokokawa for their assistance in collecting these samples. For GRADIENTS we would like to thank the captain and crew of the R/V Marcus G. Langseth (2017) and R/V Kilo Moana (2019) for their assistance in collecting these samples. For the Southern Oceans HEOBI cruise we acknowledge the use of the CSIRO Marine National Facility (https://ror.org/01mae9353) in undertaking this research. We would like to thank the captain and crew of the R/V Investigator, and the support of an Antarctic Science International Bursary awarded to Swan L. S. Sow (University of Tasmania) for assistance in collecting these samples. For the P15S cruise we acknowledge the use of the CSIRO Marine National Facility (https://ror.org/01mae9353) in undertaking this research. We would like to thank the captain and crew of the R/V Investigator for their assistance in collecting these samples. For the INS2017 cruise we acknowledge the use of the CSIRO Marine National Facility (https://ror.org/01mae9353) in undertaking this research. We would like to thank the captain and crew of the R/V Investigator and for their assistance in collecting these samples and the funding support of an Antarctic Science International Bursary awarded to Swan L. S. Sow (University of Tasmania). For the K-AXIS cruise we would like to thank the captain and crew of the R/V Aurora Australis and the Science Technical Support Team of the Australian Antarctic Division for their assistance in collecting these samples and the funding support of an Antarctic Science International Bursary awarded to Swan L. S. Sow (University of Tasmania). For the I9N and I8S cruises we would like to thank the captain and crew of the R/V Roger Revelle, UCSB at-sea team members Chantal Swan and Dave Menzies and especially the trace metals teams from U.Hawaii (Chris Measures Lab) and Florida State Univ. (Bill Landing lab) for their assistance in collecting these samples. For both P16N and P16S we would like to thank the captain and crew of the R/V Thomas G. Thompson and R/V Roger Revelle for their assistance in collecting these samples, as well as Chief Scientists Christopher L. Sabine (P16N), Richard A. Feely (P16N), and Bernadette M. Sloyan (P16S). Finally, for POTATOE, we would like to thank the captain and crew of the RVIB Nathaniel B Palmer for their assistance in collecting these samples.

## Author Contributions

Andrew P Rees and Glen Tarran supplied samples from AMT from which DNA was extracted and supplied to the Fuhrman lab by Mary R. Gradoville, Jonathan P Zehr, and Kendra Turk-Kubo. Katja Metfies, Matthias Wietz, and Christina Bienhold supplied amplicon data for FRAM. Katja Metfies, Thomas Mock, Sinhué Torres-Valdés, Sarah Lena Eggers supplied amplicon data from MOSAiC. Mary R. Gradoville, Jonathan P Zehr, Kendra Turk-Kubo, and E. Virginia Armbrust supplied extracted DNA from SCOPE-Gradients. Levente Bodrossy, Jodie Van De Kamp, Mark Brown, Swan L. S. Sow supplied extracted DNA from IND-2017, KAXIS, HEOBI, and P15S. Wade Jeffrey and Joseph Moss supplied samples from POTATOE. Paul Berube and Steven Biller supplied extracted DNA from BioGEOTRACES. GO-SHIP I08S, I09N, P16N, and P16S samples were supplied by Craig Carlson and Elisa Halewood. Jesse McNichol performed DNA extractions for I09N, I08S, P16N, P16S, and POTATOE. Swan L. S. Sow and Jodie van de Kamp performed DNA extractions from HEOBI, K-AXIS, P15S, and IND-2017, while DNA extractions from the SCOPE-Gradients cruises were performed by Mary R. Gradoville. Jesse McNichol performed PCR and library preparation for all samples with the exception of FRAM/MOSAiC with assistance from Yubin Raut and Bruce Yanpui Chan. Jesse McNichol and Nathan L R Williams developed the bioinformatic pipeline to process samples and performed bioinformatics. Nathan L R Williams collected and organized sample metadata. Nathan L R Williams, Jesse McNichol, Yubin Raut, and Jed Fuhrman wrote the manuscript, and all authors were involved in editing the manuscript.

## References

1. Woese, C. R., Kandler, O. & Wheelis, M. L. Towards a natural system of organisms: proposal for the domains Archaea, Bacteria, and Eucarya. Proc. Natl. Acad. Sci. U. S. A. 87, 4576–4579 (1990).

2. Azam, F. et al. The Ecological Role of Water-Column Microbes in the Sea. Mar. Ecol. Prog. Ser. 10, 257–263 (1983).

3. Falkowski, P. G., Fenchel, T. & Delong, E. F. The Microbial Engines That Drive Earth’s Biogeochemical Cycles. Science 320, 1034–1039 (2008).

4. Steele, J. H. The Structure of Marine Ecosystems. (Harvard University Press, 2013). doi:10.4159/harvard.9780674592513.

5. Worden, A. Z. et al. Rethinking the marine carbon cycle: Factoring in the multifarious lifestyles of microbes. Science 347, 1257594 (2015).

6. Barton, A. D. et al. The biogeography of marine plankton traits. Ecol. Lett. 16, 522–534 (2013).

7. Hanson, C. A., Fuhrman, J. A., Horner-Devine, M. C. & Martiny, J. B. H. Beyond biogeographic patterns: processes shaping the microbial landscape. Nat. Rev. Microbiol. 10, 497–506 (2012).

8. Chénard, C. et al. Temporal and spatial dynamics of Bacteria, Archaea and protists in equatorial coastal waters. Sci. Rep. 9, 16390 (2019).

9. Brown, M. V. et al. Microbial community structure in the North Pacific ocean. ISME J. 3, 1374– 1386 (2009).

10. Yeh, Y.-C. & Fuhrman, J. A. Contrasting diversity patterns of prokaryotes and protists over time and depth at the San-Pedro Ocean Time series. ISME Commun. 2, 36 (2022).

11. Chénard, C. et al. Temporal and spatial dynamics of Bacteria, Archaea and protists in equatorial coastal waters. Sci. Rep. 9, 16390 (2019).

12. Yeh, Y.-C. & Fuhrman, J. A. Effects of phytoplankton, viral communities, and warming on free-living and particle-associated marine prokaryotic community structure. Nat. Commun. 13, 7905 (2022).

13. Pesant, S. et al. Open science resources for the discovery and analysis of Tara Oceans data. Sci. Data 2, 150023 (2015).

14. Duarte, C. M. Seafaring in the 21St Century: The Malaspina 2010 Circumnavigation Expedition. Limnol. Oceanogr. Bull. 24, 11–14 (2015).

15. Alam, I. et al. KAUST Metagenomic Analysis Platform (KMAP), enabling access to massive analytics of re-annotated metagenomic data. Sci. Rep. 11, 11511 (2021).

16. Anderson, R. F., Mawji, E., Cutter, G. A., Measures, C. I. & Jeandel, C. GEOTRACES: Changing the Way We Explore Ocean Chemistry. Oceanography 27, 50–61 (2014).

17. Clayton, S. et al. Bio-GO-SHIP: The Time Is Right to Establish Global Repeat Sections of Ocean Biology. Front. Mar. Sci. 8, (2022).

18. Kopf, A. et al. The ocean sampling day consortium. GigaScience 4, 27 (2015).

19. Thompson, L. R. et al. A communal catalogue reveals Earth’s multiscale microbial diversity. Nature 551, 457–463 (2017).

20. Parada, A. E., Needham, D. M. & Fuhrman, J. A. Every base matters: assessing small subunit rRNA primers for marine microbiomes with mock communities, time series and global field samples. Environ. Microbiol. 18, 1403–1414 (2016).

21. Bolyen, E. et al. Reproducible, interactive, scalable and extensible microbiome data science using QIIME 2. Nat. Biotechnol. 37, 852–857 (2019).

22. Callahan, B. J. et al. DADA2: High-resolution sample inference from Illumina amplicon data. Nat. Methods 13, 581–583 (2016).

23. Yeh, Y. et al. Comprehensive single-PCR 16S and 18S rRNA community analysis validated with mock communities, and estimation of sequencing bias against 18S. Environ. Microbiol. 23, 3240–3250 (2021).

24. McNichol, J., Berube, P. M., Biller, S. J. & Fuhrman, J. A. Evaluating and Improving Small Subunit rRNA PCR Primer Coverage for Bacteria, Archaea, and Eukaryotes Using Metagenomes from Global Ocean Surveys. mSystems 6, e00565–21 (2021).

25. Baldwin, A., Moss, J., Pakulski, J., Joux, F. & Jeffrey, W. Microbial diversity in a Pacific Ocean transect from the Arctic to Antarctic circles. Aquat. Microb. Ecol. 41, 91–102 (2005).

26. Moss, J. A., Henriksson, N. L., Dean, P. J., Snyder, R. A. & Jeffrey, W. H. Oceanic Microplankton Do Not Adhere to the Latitudinal Diversity Gradient. Microb. Ecol. 79, 511–515 (2020).

27. Lee, Z. et al. Euphotic zone depth: Its derivation and implication to ocean-color remote sensing. J. Geophys. Res. Oceans 112, (2007).

28. Kirk, J. T. O. *Light and Photosynthesis in Aquatic Ecosystems*. (Cambridge University Press, Cambridge, 2010). doi:10.1017/CBO9781139168212.

29. Varaljay, V. A. et al. Single-taxon field measurements of bacterial gene regulation controlling DMSP fate. ISME J. 9, 1677–1686 (2015).

30. Biller, S. J. et al. Marine microbial metagenomes sampled across space and time. Sci. Data 5, 180176 (2018).

31. Gradoville, M. R. et al. Latitudinal constraints on the abundance and activity of the cyanobacterium UCYN-A and other marine diazotrophs in the North Pacific. Limnol. Oceanogr. 65, 1858–1875 (2020).

32. Brown, M. V. et al. Systematic, continental scale temporal monitoring of marine pelagic microbiota by the Australian Marine Microbial Biodiversity Initiative. Sci. Data 5, 180130 (2018).

33. Raes, E. J., Bodrossy, L., van de Kamp, J., Bissett, A. & Waite, A. M. Marine bacterial richness increases towards higher latitudes in the eastern Indian Ocean. Limnol. Oceanogr. Lett. 3, 10– 19 (2018).

34. Sow, S. L. S. et al. Biogeography of Southern Ocean prokaryotes: a comparison of the Indian and Pacific sectors. Environ. Microbiol. 24, 2449–2466 (2022).

35. Boström, K. H., Simu, K., Hagström, Å. & Riemann, L. Optimization of DNA extraction for quantitative marine bacterioplankton community analysis. Limnol. Oceanogr. Methods 2, 365–373 (2004).

36. Manganelli, M. et al. Major Role of Microbes in Carbon Fluxes during Austral Winter in the Southern Drake Passage. PLoS ONE 4, e6941 (2009).

37. Signori, C. N., Thomas, F., Enrich-Prast, A., Pollery, R. C. G. & Sievert, S. M. Microbial diversity and community structure across environmental gradients in Bransfield Strait, Western Antarctic Peninsula. Front. Microbiol. 5, (2014).

38. Lila Koumandou, V., Nisbet, R. E. R., Barbrook, A. C. & Howe, C. J. Dinoflagellate chloroplasts – where have all the genes gone? Trends Genet. 20, 261–267 (2004).

39. Sanchez-Garcia, S., Wang, H. & Wagner-Döbler, I. The microbiome of the dinoflagellate Prorocentrum cordatum in laboratory culture and its changes at higher temperatures. Front. Microbiol. 13, (2022).

40. 40. Vaulot, D. pr2-primers version 2.0.0. (2023).

41. Quast, C. et al. The SILVA ribosomal RNA gene database project: improved data processing and web-based tools. Nucleic Acids Res. 41, D590–D596 (2012).

42. Kelly, L., Huang, K. H., Ding, H. & Chisholm, S. W. ProPortal: a resource for integrated systems biology of Prochlorococcus and its phage. Nucleic Acids Res. 40, D632–D640 (2012).

43. 43. R Core Team. A Language and Environment for Statistical Computing. (2021).

44. Dixon, P. VEGAN, A Package of R Functions for Community Ecology. J. Veg. Sci. 14, 927–930 (2003).

45. Wickham, H. et al. Welcome to the Tidyverse. J. Open Source Softw. 4, 1686 (2019).

46. Kassambara, A. ggpubr: “ggplot2” Based Publication Ready Plots. (2023).

47. Pedersen, T. L. patchwork: The Composer of Plots. (2024).

48. Ashkezari, M. D. et al. Simons Collaborative Marine Atlas Project (Simons CMAP): An open-source portal to share, visualize, and analyze ocean data. Limnol. Oceanogr. Methods 19, 488–496 (2021).

49. Sunagawa, S. et al. Tara Oceans: towards global ocean ecosystems biology. Nat. Rev. Microbiol. 18, 428–445 (2020).

50. Raina, J.-B. et al. Chemotaxis shapes the microscale organization of the ocean’s microbiome. Nature 605, 132–138 (2022).

51. Azam, F. & Malfatti, F. Microbial structuring of marine ecosystems. Nat. Rev. Microbiol. 5, 782–791 (2007).

52. Xu, S. et al. Diversity, community structure, and quantity of eukaryotic phytoplankton revealed using 18S rRNA and plastid 16S rRNA genes and pigment markers: a case study of the Pearl River Estuary. Mar. Life Sci. Technol. 5, 415–430 (2023).

53. Needham, D. M. & Fuhrman, J. A. Pronounced daily succession of phytoplankton, archaea and bacteria following a spring bloom. Nat. Microbiol. 1, 16005 (2016).

54. White, M. M. & McLaren, I. A. Copepod development rates in relation to genome size and 18S rDNA copy number. Genome 43, 750–755 (2000).

55. Jones-Kellett, A. E. et al. Amplicon sequencing with internal standards yields accurate picocyanobacteria cell abundances as validated with flow cytometry. ISME Commun. 4, ycae115 (2024).

