## Supplementary Figure 1 for "Characterizing organisms from three domains of life with universal primers from throughout the global ocean"

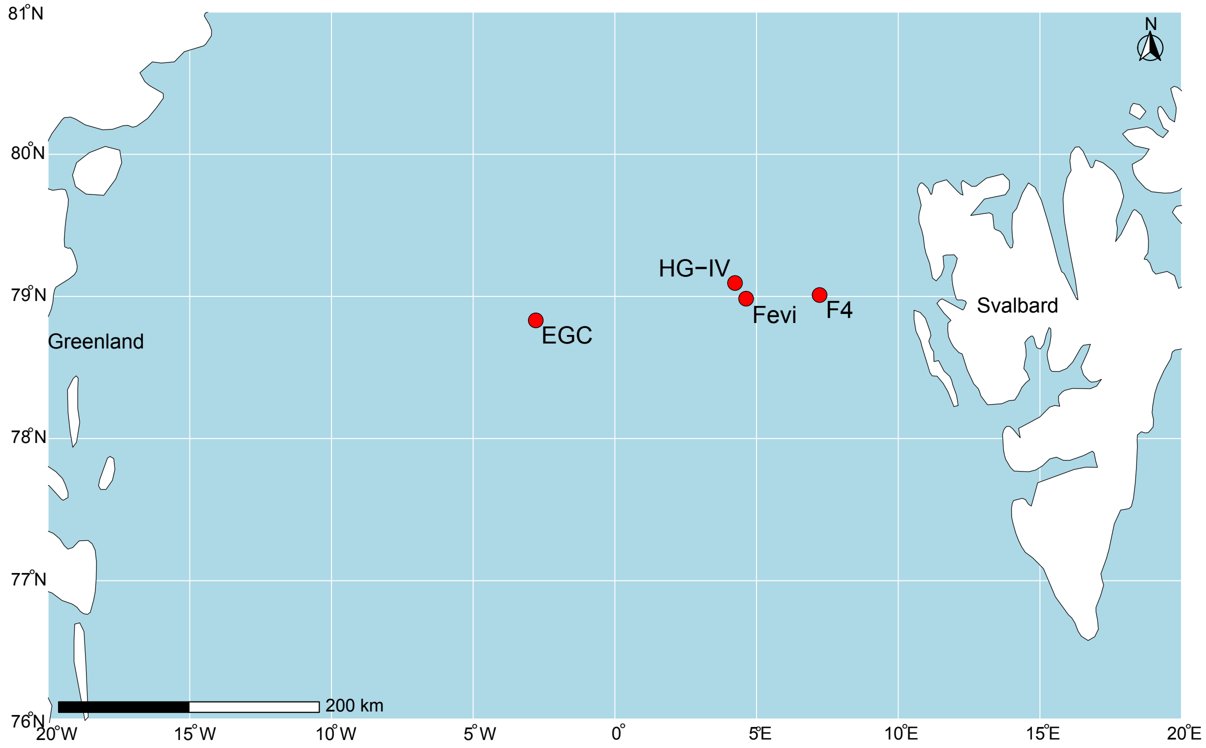


**Supplementary Figure 1** Map of FRAM sampling locations.

**Supplementary Table 1.** Correction factors for both 16S and 18S sequences. The Southern Oceans and Gradients Bioanalyzer data are in units pmole/L.

| SEQUENCING RUN | 16S BIOANLYZER | 18S BIOALYZER | 16S FRACTION BIOANALYZER | 18S FRACTION-BIOANALYZER | 16S SEQUENCES | 18S SEQUENCES | 16S FRACTION SEQUENCES | 18S FRACTION SEQUENCES | 16S CORRECTION FACTOR | 18S CORRECTION FACTOR |
|---|---|---|---|---|---|---|---|---|---|---|
| **SOUTHERN_OCEANS** | 1340.0 | 335.1 | 0.800 | 0.199 | 29501073 | 3462334 | 0.894 | 0.105 | 0.89 | 1.90 |
| **GRADIENTS** | 3292.0 | 638.5 | 0.837 | 0.162 | 101174114 | 7014632 | 0.935 | 0.064 | 0.90 | 2.51 |
| **P16N_S** | 6162.1 | 2182.8 | 0.738 | 0.261 | 42116230 | 3082200 | 0.931 | 0.068 | 0.79 | 3.84 |
| **GA03_GP13** | 2052.0 | 434.0 | 0.82 | 0.17 | 24433763 | 734205 | 0.970 | 0.029 | 0.85 | 5.98 |
| **I8_I9** | 6162.1 | 2185.2 | 0.738 | 0.261 | 40226353 | 224586 | 0.947 | 0.052 | 0.78 | 4.95 |
| **GA02_GA10** | 124014.1 | 34444.1 | 0.782 | 0.217 | 21661905 | 678869 | 0.969 | 0.030 | 0.81 | 7.15 |
| **AMT_19_20_POTATOE** | 5969.3 | 1361.0 | 0.814 | 0.18 | 11250567 | 226122 | 0.980 | 0.019 | 0.83 | 9.42 |
| **FRAM_MOSAIC** | NA | NA | NA | NA | 24349430 | 2035473 | 0.922 | 0.077 | 0.89 | 1.90 |

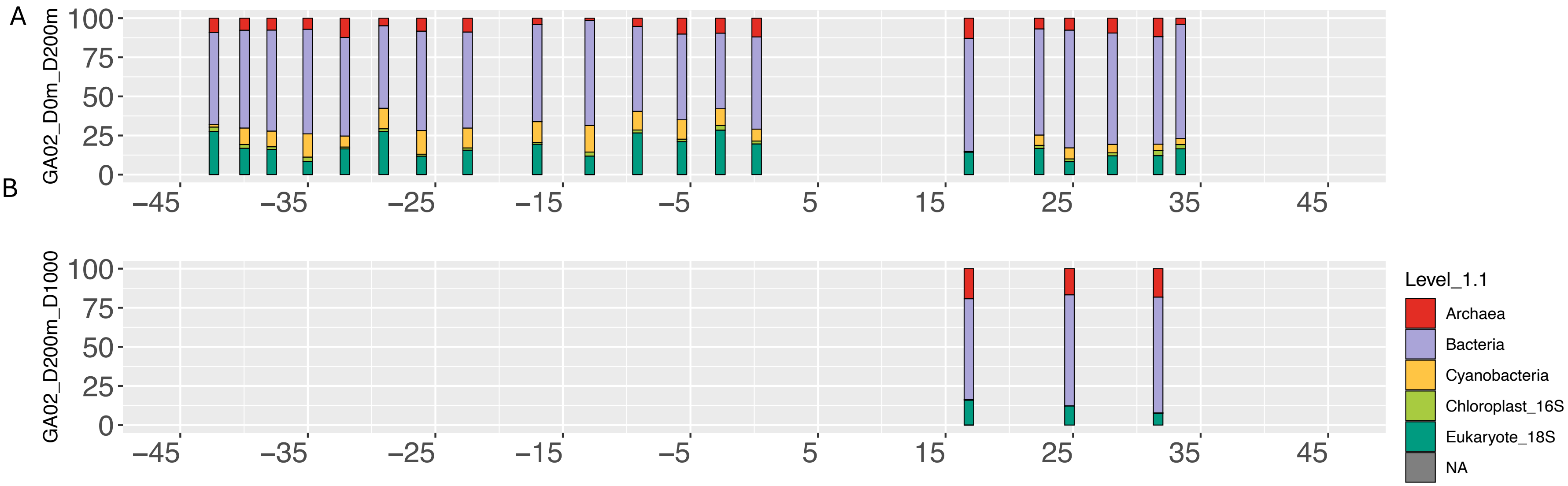


**Supplementary Figure 2.** Relative abundance of marine microbes from the GA02 transect from the surface – 200 m of the ocean, and 200 m - 1000 m, partitioned between Archaea, Bacteria, Cyanobacteria, Chloroplast 16S, and Eukaryotic 18S. Relative abundance (y axis) is shown at increasing latitudes (x axis) from within the Atlantic Ocean (Cruise GA02). A is top 200m, B is 200-1000m depths.

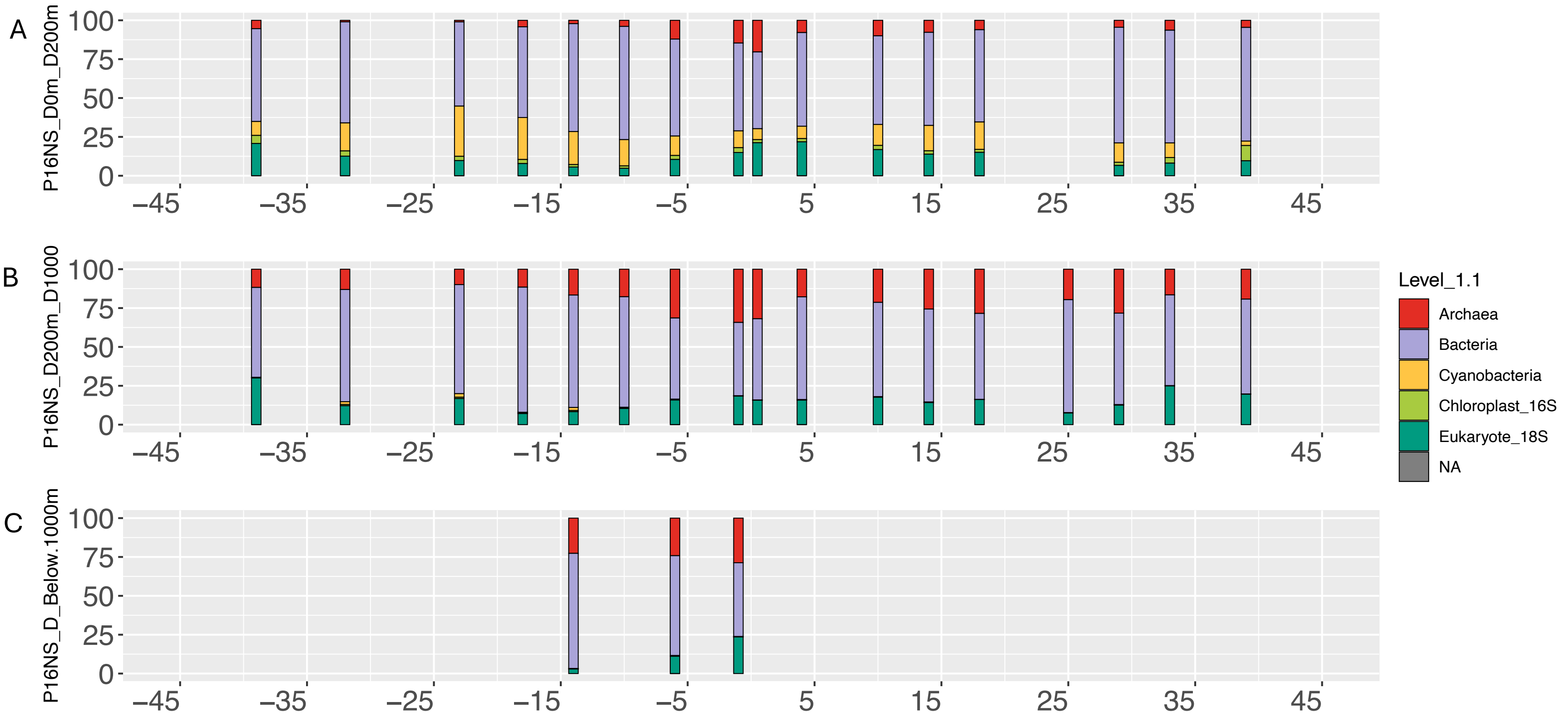


**Supplementary Figure 3.** Relative abundance of marine microbes from the P16N cruise between the surface – 200 m of the ocean, 200 m - 1000 m, and then below 1000 m, partitioned between Archaea, Bacteria, Cyanobacteria, Chloroplast 16S, and Eukaryotic 18S. Relative abundance (y axis) is shown at increasing latitudes (x axis) from within the Pacific Ocean (Cruise P16N and P16S). A is top 200m, B is 200-1000m, C is below 1000m.

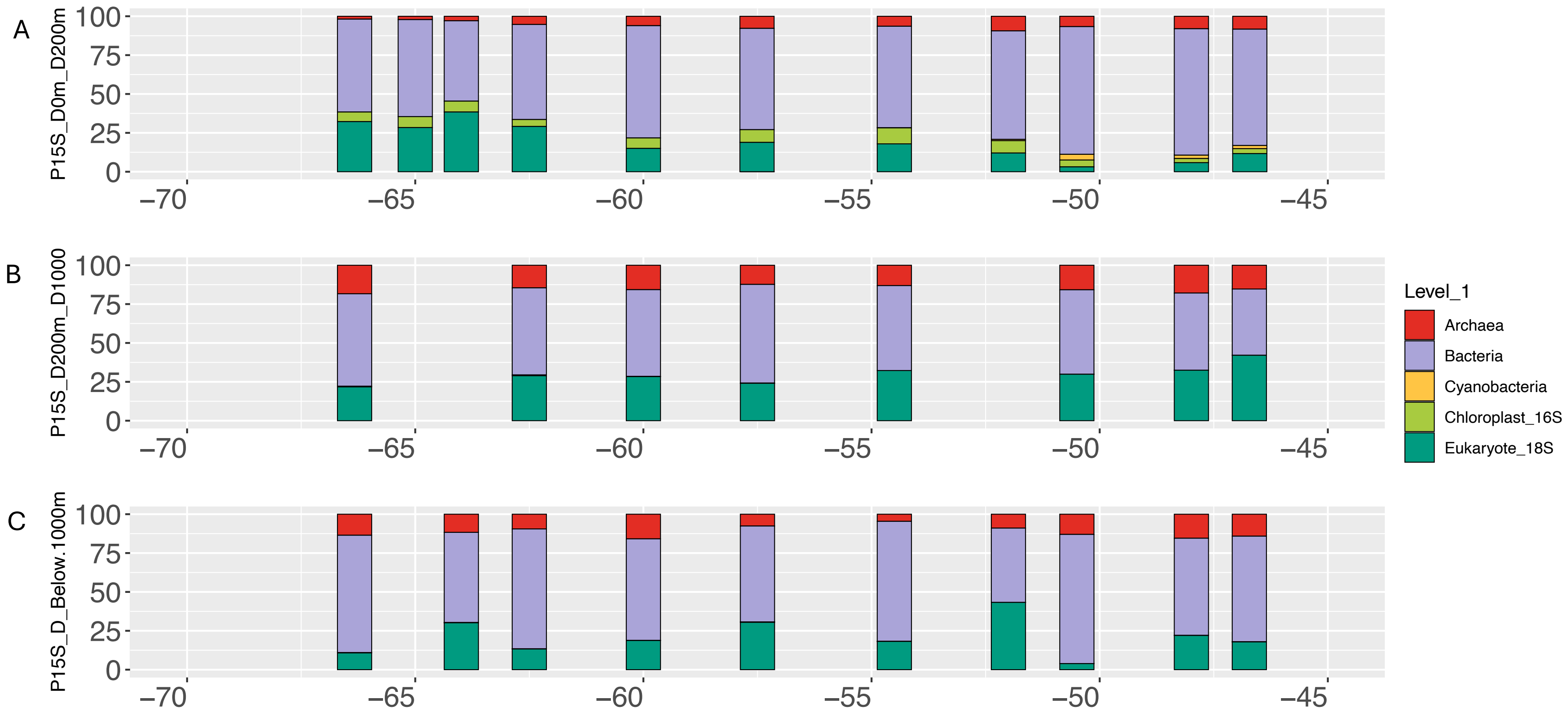


**Supplementary Figure 4.** Relative abundance of marine microbes from the P15S transect between the surface – 200 m of the ocean, 200 m - 1000 m, and then below 1000 m, partitioned between Archaea, Bacteria, Cyanobacteria, Chloroplast 16S, and Eukaryotic 18S. Relative abundance (y axis) is shown at increasing latitudes (x axis) from within the Southern Ocean (Cruise P15S). A is top 200m, B is 200-1000m, C is below 1000m.

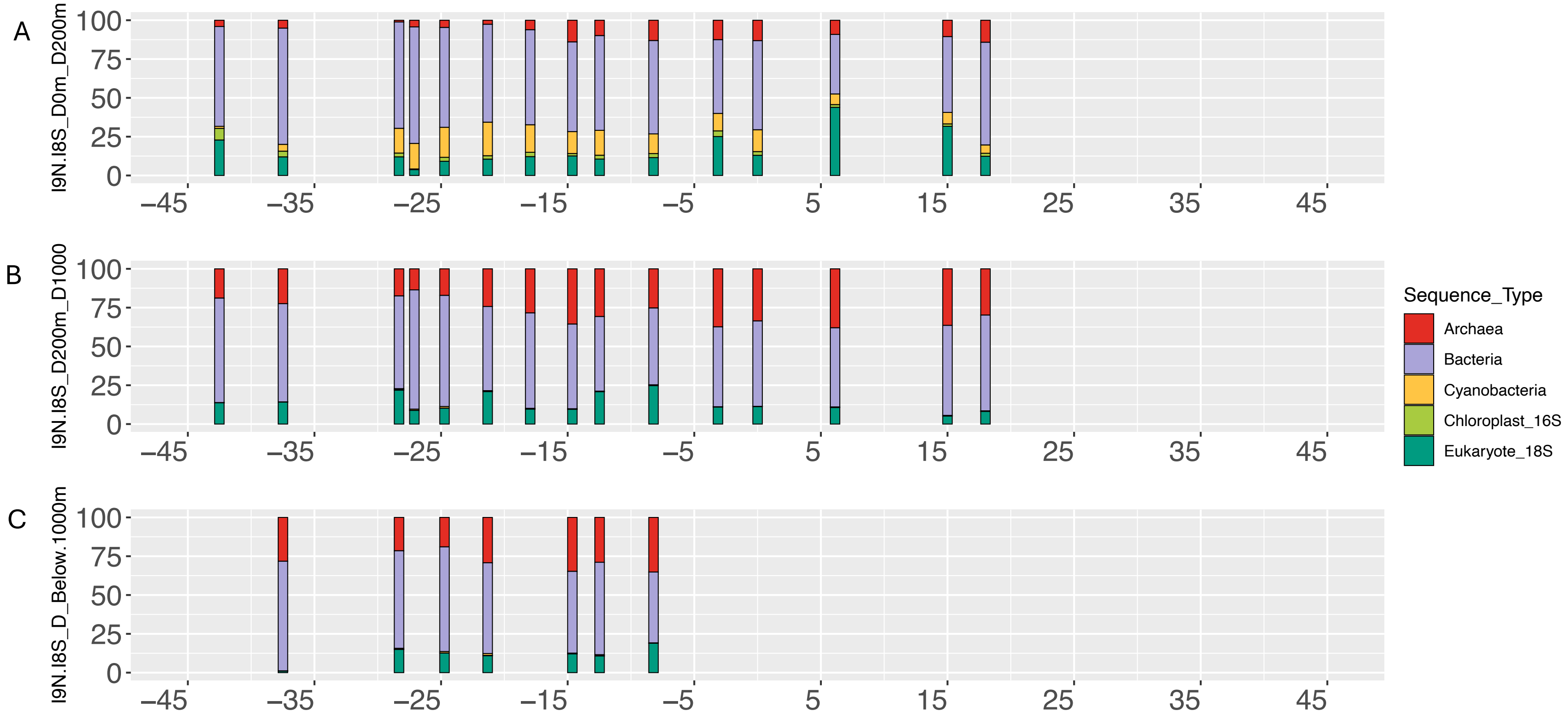


**Supplementary Figure 5.** Relative abundance of marine microbes from the I9N and I8S transects between the surface – 200 m of the ocean, 200 m - 1000 m, and then below 1000 m partitioned between Archaea, Bacteria, Cyanobacteria, Chloroplast 16S, and Eukaryotic 18S. Relative abundance (y axis) is shown at increasing latitudes (x axis) from within the Indian Ocean (Cruise IO9N and IO8S). A is top 200m, B is 200-1000m, C is below 1000m.
